# A Non-Classical Neuroactive Steroid Exhibiting Potent, Efficacious GABA_A_ Receptor Agonism and NMDA Receptor Inhibition

**DOI:** 10.64898/2026.04.06.716659

**Authors:** Hong-Jin Shu, Yuanjian Xu, Mingxing Qian, Ann Benz, Carla M. Yuede, Douglas F. Covey, Charles F. Zorumski, Steven Mennerick

## Abstract

Neuroactive steroids modulate GABA_A_ and NMDA receptors allosterically, typically requiring specific structural features for their activity. In this study, we characterize YX84, a novel neuroactive steroid bearing a 3β sulfate and *p-*trifluoroacetylbenzyl alcohol attached in an ether linkage to a hydroxyl group at steroid carbon 17. This compound and similar analogues exhibit an atypical pharmacological profile, with three distinct actions at GABA_A_ receptors. First, YX84 is a full agonist, with EC_50_ near 1 µM and comparable efficacy to GABA at GABA_A_ receptors in native hippocampal neurons. It presents as a full agonist relative to GABA at α4/δ subunit-containing receptors. Second, YX84 acts as a slow-onset, potent positive allosteric modulator (PAM) of GABA_A_ receptors at concentrations below those that gate a response. Finally, YX84 exhibits rapid desensitizing and/or blocking kinetics; voltage dependence is consistent with a contribution of channel block. Structure– activity relationship analyses reveal that both functional groups are essential for gating activity, while classical requirements such as carbon 3 hydroxyl stereoselectivity and carbon 5 reduction are dispensable. YX84 also modestly inhibits NMDA receptor currents, suggesting weak negative allosteric modulation. Behavioral assays show that intraperitoneal administration of YX84 (30 mg/kg) does not impair sensorimotor function, unlike allopregnanolone. These findings identify YX84 as a structurally distinct neuroactive steroid with dual receptor activity and favorable behavioral tolerability, offering a promising scaffold for therapeutic development targeting excitatory/inhibitory imbalance in neuropsychiatric disorders if pharmacokinetic considerations can be overcome.

## 1. Introduction

Neuroactive steroids are emerging as a promising class of therapeutics for managing brain disorders, including epilepsy and mood disorders (Deligiannidis and Meltzer-Brody, 2025; Zorumski et al., 2025; Bortolato and Pinna, 2026). FDA-approved compounds function as positive allosteric modulators (PAMs) of GABA_A_ receptors at low concentrations (Martinez Botella et al., 2015). Like other PAMs, such as propofol and barbiturates, these neuroactive steroids can directly activate GABA_A_ receptors at high concentrations, exhibiting low-efficacy, slow gating kinetics in the absence of orthosteric agonists (Shu et al., 2004). A distinct subset of neuroactive steroids, characterized by a sulfate group at C3 and exemplified by pregnenolone sulfate, negatively modulates GABA_A_ receptor function and variably affects NMDAR function (Park-Chung et al., 1999; Weaver et al., 2000). Crucially, potent GABA_A_R PAM activity is diastereoselective at C3 (Covey et al., 2001). Sulfation at C3 represents a regulatory point that inverts GABA_A_ receptor modulation from PAM to negative allosteric modulation (NAM) and imparts NMDAR PAM or NAM activity (Park-Chung et al., 1997).

During our recent exploration of neuroactive steroid photoaffinity labeling agents targeting GABA_A_ and NMDA receptors, we identified several compounds with unique structural features that violate the structural norms above and appeared to exert allosteric effects on GABA_A_ receptors and NMDA receptors (Qian et al., 2023). Pursuing this work further, here we investigate the structure-activity profile and pharmacological properties at GABA_A_ receptors and an initial investigation of NMDA receptor interactions. We uncovered an atypical constellation of GABA_A_ receptor actions that include efficacious direct agonism, positive modulation, and apparent desensitization mediated by atypical structure-activity features. The compound highlighted in this report, YX84, emerged as a lead tool to explore this unusual pharmacology.

Neuroactive steroids with improved pharmacological profiles may need to incorporate structural modifications that depart from classical neurosteroid scaffolds. In this context, compounds represented by YX84 are intriguing because they incorporate features not typically associated with GABA_A_ receptor potentiators or agonists. Accordingly, these compounds appear to engage GABA_A_ receptors through a pharmacological profile distinct from that of typical neurosteroids or barbiturates and also exhibit measurable negative allosteric effects at NMDA receptors, which could offer fortuitous therapeutic synergy (Mennerick et al., 2001; Ziolkowski et al., 2021; Abramova et al., 2023).

Collectively, our pharmacological exploration expands understanding of structure-activity relationships for neuroactive steroids on GABA_A_ and NMDA receptors. If pharmacokinetic considerations can be overcome, these compounds represent a new class of modulators with potentially fortuitous dual actions at GABA_A_ receptors and at NMDA receptors. Our research contributes to growing understanding by revealing novel mechanisms of action and potential therapeutic applications of neuroactive steroids.

## 2. Methods

### 2.1 Cell Culture

Primary hippocampal neurons were cultured from postnatal day 0–2 Sprague-Dawley rats of either sex, following protocols approved by the Washington University Institutional Animal Care and Use Committee. Dissections were performed in ice-cold medium, with cells dissociated using papain digestion followed by trituration. Neurons were plated onto poly-D-lysine-coated glass coverslips at a density of ∼50,000 cells/cm² and maintained in Neurobasal-A medium supplemented with B27, GlutaMAX, and penicillin-streptomycin. Cultures were maintained at 37°C in a humidified 5% CO₂ incubator and used for recordings between 5–8 days in vitro.

### 2.2 Heterologous Expression in N2A Cells

To assess subunit-specific effects of YX84, we performed electrophysiological recordings in acutely transfected N2A cells. Cells were plated on poly-D-lysine-coated coverslips and transfected 24–48 hours prior to recording using Lipofectamine 2000 (Thermo Fisher) with plasmids encoding GABA_A_ receptor subunits of interest (e.g., α1β2γ2 or α4β2δ). A plasmid encoding GFP was co-transfected to identify successfully transfected cells. Recordings were performed under voltage-clamp conditions using the same internal and external solutions as described for hippocampal neurons. Transfected cells were selected based on GFP fluorescence and morphology, and recordings were conducted 24–48 hours post-transfection.

### 2.3 Electrophysiology

We performed whole-cell voltage-clamp recordings at room temperature using an Axopatch 200B amplifier (Molecular Devices). Patch pipettes (3–5 MΩ) were pulled from borosilicate glass and filled with an internal solution containing (in mM): 140 CsCl, 10 HEPES, 5 EGTA, 2 MgATP, and 0.3 NaGTP (pH 7.3, osmolarity ∼290 mOsm). Neurons were continuously perfused with external solution containing (in mM): 140 NaCl, 3 KCl, 2 CaCl₂, 1 MgCl₂, 10 HEPES, 0.025 D-APV, 0.001 NBQX, and 10 glucose (pH 7.4). Drugs were applied through a common port via a gravity-fed, multi-barrel local perfusion system. Currents were filtered at 2 kHz and digitized at 10 kHz using pClamp software. Series resistance was monitored throughout.

### 2.4 Sensorimotor Behavioral Testing

Mice were habituated to the testing room for 30–60 minutes prior to testing. Sensorimotor function was assessed using a standardized battery of tests conducted by the Washington University Animal Behavior Core, as previously described (Chen et al., 2021). Testing was conducted during the light phase of the light/dark cycle by a female experimenter blind to the drug treatment group. The battery was performed in succession within 10 min following drug administration and included assessments of walking initiation, ledge balance, platform balance, pole ascent and descent, and inverted screen grip strength. Maximum time on each test was 60s, except for the Pole test which was 120s. Mice were tested consecutively on each apparatus, with two trials per test conducted back-to-back. The average of the two trials was used for analysis. Apparatuses were cleaned with 70% ethanol or 0.02% chlorhexidine (Nolvasan) between animals only if soiled. Testing order was fixed across animals to ensure consistency. Ordering of drug conditions (below) was randomized.

### 2.5 In Vivo Drug Administration

Immediately prior to behavioral testing, mice received intraperitoneal injections of either 15 mg/kg allopregnanolone, 30 mg/kg YX84, or vehicle control (20% (2-hydroxypropyl)- β-cyclodextrin in saline). The choice of allopregnanolone dose was informed by previous studies by our group suggesting that this dose would represent the threshold of sensorimotor effects (Salvatore et al., 2024; Lambert et al., 2025). Choice of YX84 dose was based on potent *in vitro* pharmacodynamics described below, paired with the assumption that pharmacokinetics may be less favorable than for allopregnanolone. All injections were administered in a volume appropriate for the animal’s weight, and behavioral testing commenced within 10 minutes of injection to capture acute drug effects. Behavioral testing was completed in 15 minutes. Weight was recorded for each animal near the time of testing to account for potential influences on task performance.

### 2.6 Reagents

YX84 and its analogues were synthesized in-house. Synthetic details are reported in Supplemental Information. Comparators were obtained from commercial sources when available. Allopregnanolone and 17-phenyl-(3α,5α)-androst-16-en-3-ol (17PA) were purchased from Tocris Bioscience. GABA, picrotoxin, bicuculline, gabazine, pentobarbital, and (2-hydroxypropyl)-β-cyclodextrin were obtained from Sigma-Aldrich. Stock solutions for *in vitro* experiments were prepared in DMSO or water depending on solubility and diluted into external solution immediately before use. Final DMSO concentrations did not exceed 0.1%.

### 2.7 Data Analysis

Electrophysiological data were analyzed using Clampfit (Molecular Devices). For potentiation, peak current amplitudes were measured relative to baseline and normalized to control responses where appropriate. For gating, peak responses to sub-saturating YX84 concentrations were normalized to the response of a saturating concentration.

Concentration-response curves were fit using the Hill equation: I = Iₘₐₓ × [D]ⁿ/(EC₅₀ⁿ + [D]ⁿ) where I is the response amplitude, [D] is the drug concentration, EC_50_ is the concentration producing half-maximal response, and *n* is the Hill coefficient. Behavioral data were plotted and statistically analyzed using GraphPad Prism. Statistical comparisons were made using one-way ANOVA followed by Tukey’s post hoc test, with significance defined as *p* < 0.05. Data are presented as mean ± SEM.

## 3. Results

### 3.1 GABA_A_ Receptor Gating

YX84 (5 µM; Figure 1A) gated large, peak inward currents characterized by rapid apparent desensitization in cultured hippocampal neurons (Figure 1B) at modest concentrations relative to gating by typical neurosteroids (Shu et al., 2004). The YX84 current was sensitive to 50 µM picrotoxin, confirming mediation via GABA_A_ receptors (Figure 1C). Quantification of the concentration-response relationship for gating of peak current revealed an EC₅₀ of 0.9 µM and a Hill coefficient of 3.1, indicating potency 10-20-fold greater than GABA (Shu et al., 2004) and cooperative effects of YX84 in gating (Figure 1D). At a concentration subthreshold to gating (0.1 µM), YX84 potentiated responses to 0.5 µM GABA with sustained application, consistent with positive allosteric modulation (Figure 1E, F).

**Figure 1.**
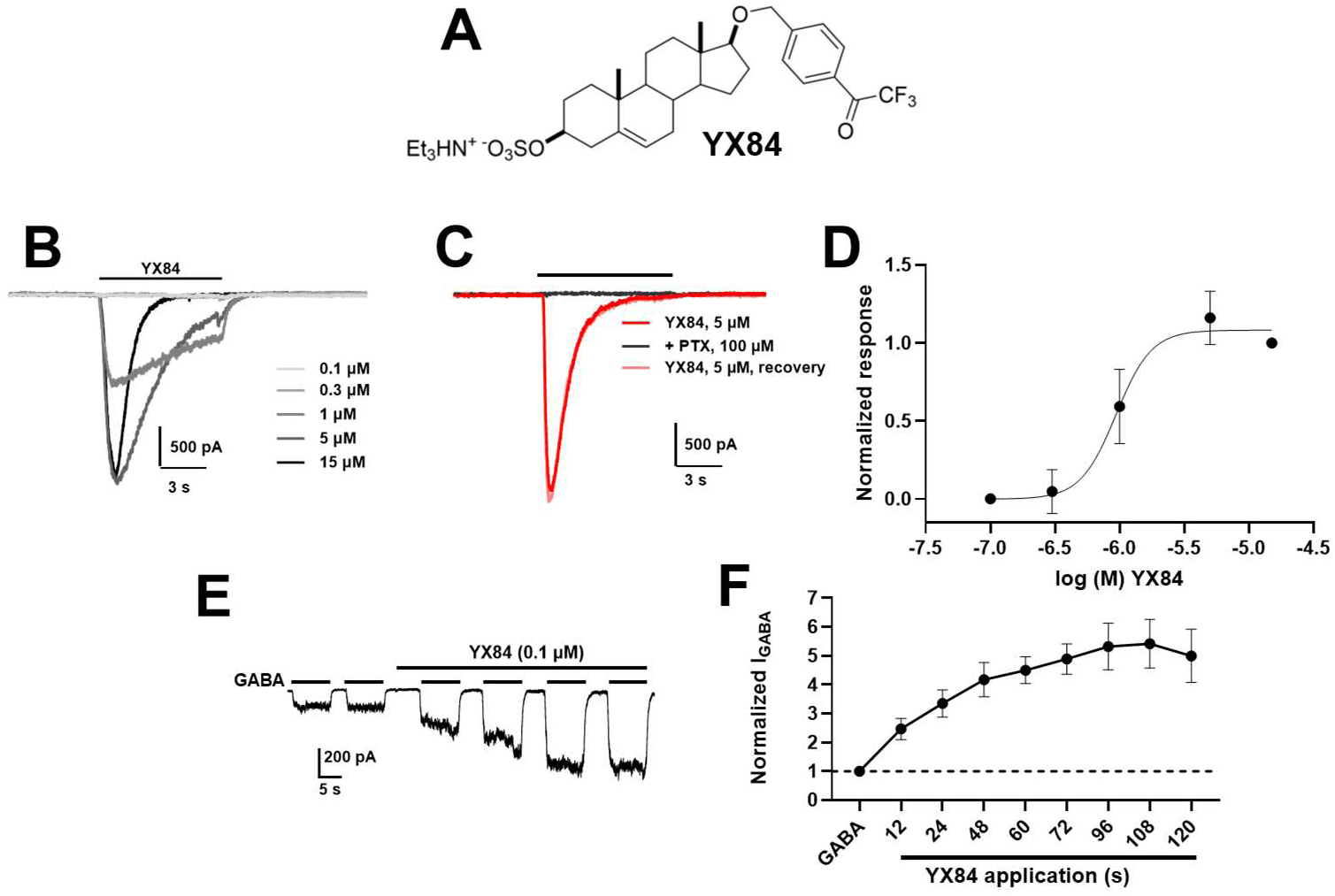
YX84 is a potent GABA_A_ receptor agonist and positive allosteric modulator of GABA_A_ receptors. **A.** Structure of YX84. Note the 3β sulfate and tri-fluoromethyl ketone at opposite ends of the steroid backbone. **B.** On a cultured hippocampal neuron, Y84 gates a concentration-dependent inward current. **C.** YX84 (5 µM)-gated currents are sensitive to the non-competitive GABA_A_ receptor antagonist picrotoxin (50 µM). **D.** Concentration-response relationship for YX84-gated current on hippocampal neurons (n = 11). Based on the fit to averaged data, the EC50 was 0.9 µM and the Hill coefficient was 3.1. **E.** At a concentration below that which gates GABA_A_ responses, YX84 (0.1 µM) exhibited a slow-onset potentiation of GABA (0.5 µM) responses. **F.** Summary of the onset time course of YX84 potentiation (n = 8).

Structure-activity relationship analysis (Figure 2) demonstrated that both the 3β sulfate and the trifluoromethyl ketone moieties were essential for receptor gating (Figure 2). Notably, activity was insensitive to stereochemistry at C3 or C5, and reduction at C5 was not required (Figure 2A), a distinct pattern from established neurosteroid structure-activity relationships (Covey et al., 2001).

**Figure 2.**
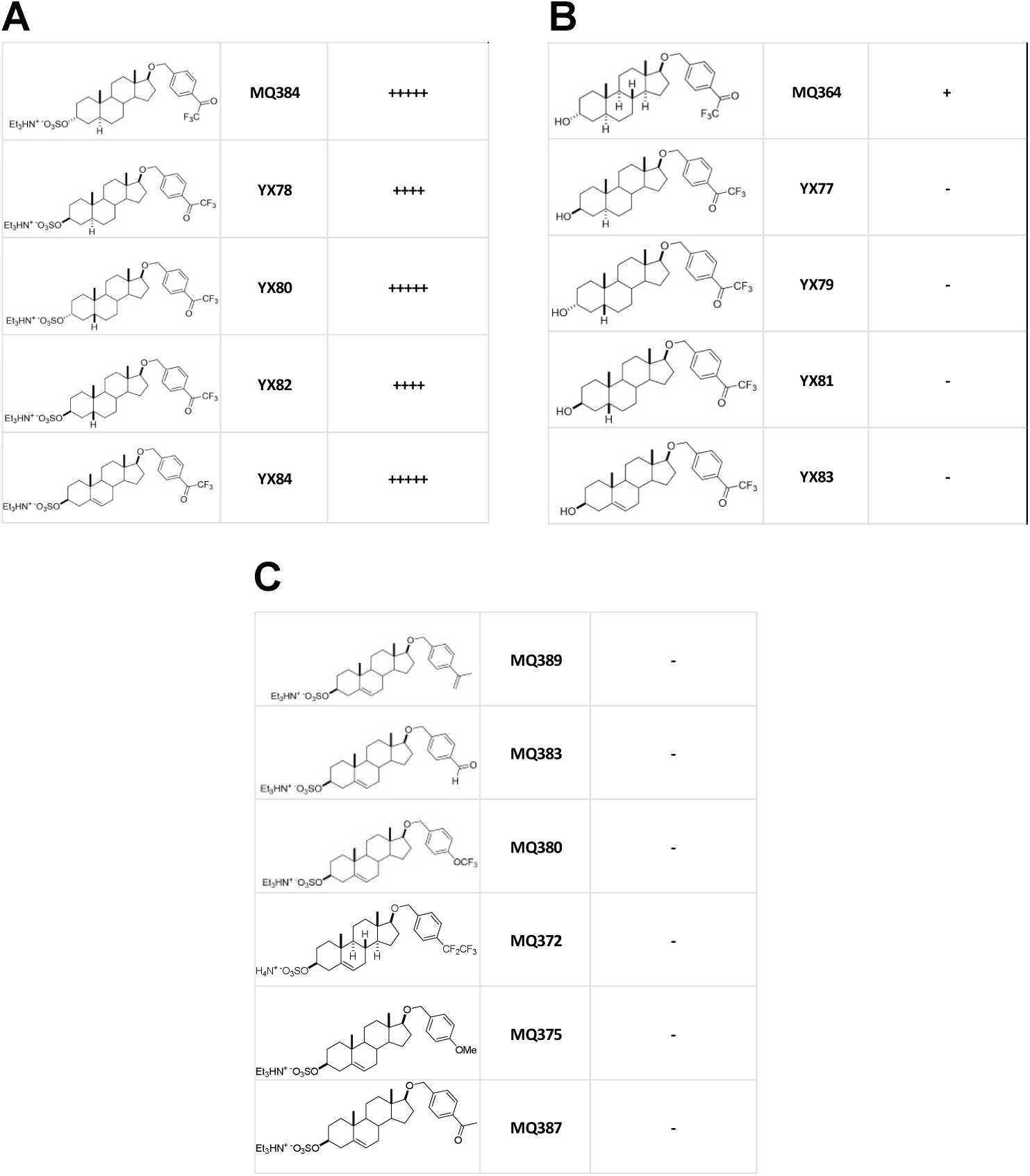
Semiquantitative summary of GABA_A_ receptor gating at a single concentration (5 µM) of various structural analogues of YX84 applied to hippocampal neurons in the absence of GABA. The left column shows structure, the middle column gives compound name, and the right column gives a relative assessment of peak current gated. No observable current was gated by compounds denotes as a ‘-‘. **A.** Variants in which stereochemistry at C3 and C5 were altered but the sulfate at C3 and the trifluoromethyl ketone off the aromatic ring were retained. **B.** Variants in which a hydroxyl group was substituted for the sulfate at C3. **C.** Substitutions for the trifluoromethyl ketone off the aromatic ring. Based on these results, we conclude that both the sulfate group at carbon 3 and the trifluoromethyl ketone on the aromatic ring at carbon 17 are required for gating. Activity is agnostic to the stereochemistry at C3.

Comparison of peak currents gated by YX84 and saturating GABA (100 µM) revealed similar amplitudes (Figure 3), suggesting that YX84 acts with equivalent efficacy to GABA at GABA_A_ receptors in hippocampal neurons under the conditions tested, which likely includes a mixture of native GABA_A_ subunit combinations.

**Figure 3.**
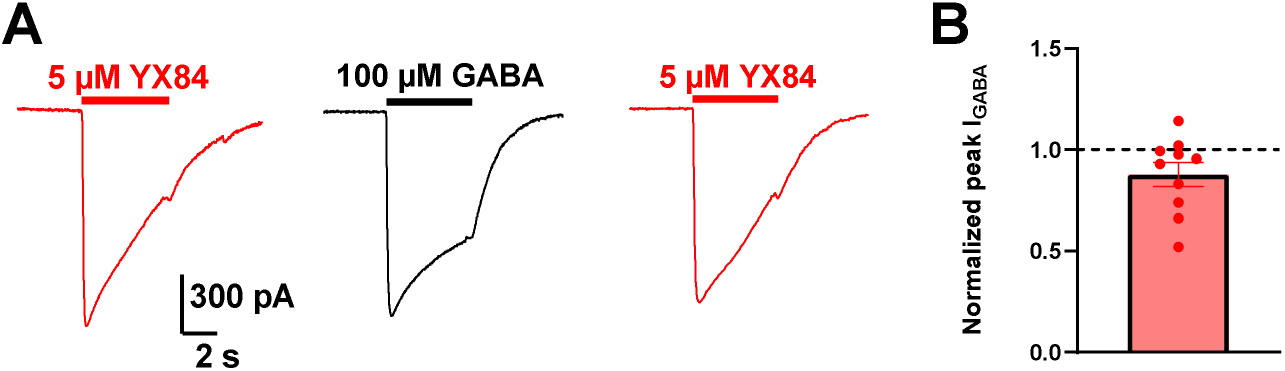
Efficacy of YX84 is equivalent to that of GABA in hippocampal neurons. **A.** We compared peak current gated by YX84 (5 µM) with that of a saturating concentration of GABA (100 µM). **B.** On average, the peak current amplitude of YX84 was similar to that of GABA (n= 10).

The rapid decrement of YX84 responses in the continued presence of YX84 could indicate agonist-dependent conformational changes in the protein (Patneau et al., 1992) or could indicate channel block by the compound, reminiscent of pregnenolone sulfate (Legesse et al., 2023). Channel block might predict voltage-dependent characteristics of the current decline, given the negative charge on the YX84 sulfate group. Consistent with weak voltage dependent channel block, we detected mild inward rectification in current-voltage relationships for YX84 (Figure 4A, C) and a faster time constant of decay during YX84 application at +50 mV relative to −70 mV (Figure 4A, D). By contrast, GABA showed a linear peak current-voltage relationship at a saturating GABA concentration (Mennerick et al., 2001; O’Toole and Jenkins, 2012)(Figure 4C), and slowed time constant of desensitization at +50 mV (Figure 4B,E).

**Figure 4.**
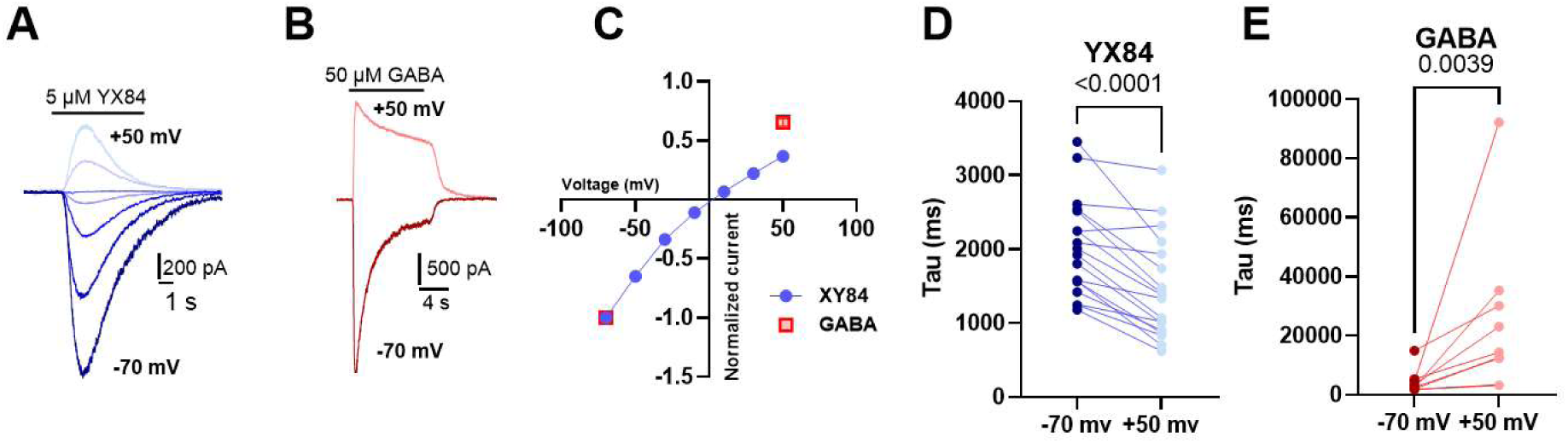
YX84 rectification consistent with channel block. **A.** Representative example of YX84 responses at varied membrane potentials (20 mV increments) from an individual hippocampal neuron. **B.** For comparison, responses to GABA in a different neuron are shown at the extreme potentials of −70 mV and +50 mV. **C.** Average peak responses, normalized to the response at −70 mV, from 22 neurons challenged with YX84 and 9 neurons with GABA. The normalized response at +50 mV differs between YX84 and GABA (p < 0.0001, Student’s *t* test), suggesting inward rectification of YX84 relative to GABA. **D, E.** Average single-exponential tau values fit to the desensitizing response at extreme voltages for YX84 (D) and for GABA (E).

Neurosteroids may have preference for GABA_A_ receptors containing a δ subunit (Brown et al., 2002; Wohlfarth et al., 2002). To test whether YX84 gating shares this preference, we compared a fixed concentration of YX84 at α4β2δ receptors and at α1β2γ2 receptors expressed in N2a cells. We used Zn^2+^ (1 µM) insensitivity to validate γ2 presence and DS2 (1 µM) sensitivity to validate δ presence (Wafford et al., 2009; Jensen et al., 2013). YX84 proved a strong agonist at both receptor subtypes but was especially efficacious at α4/δ receptors, in contrast to GABA, a partial agonist at these receptors (Brown et al., 2002; Belelli et al., 2005; Storustovu and Ebert, 2006)(Figure 5A, C).

**Figure 5.**
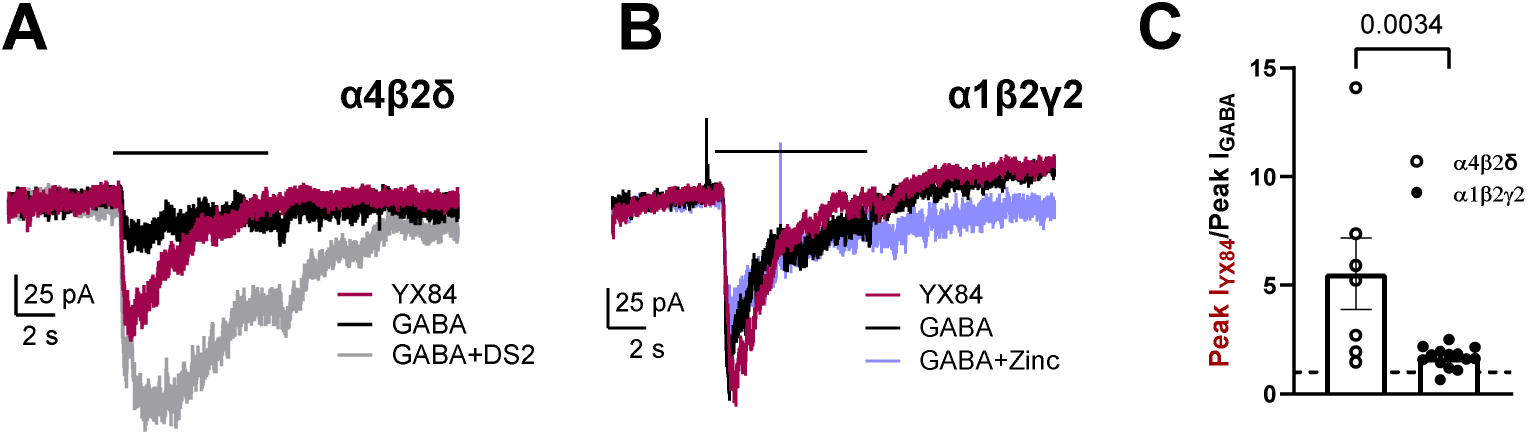
YX84 has strong efficacy at α4/δ subunit-containing receptors. N2a cells were transiently transfected with the subunits indicated in the panels. **A.** GABA (10 µM) responses (black trace) were validated for δ presence by challenge with co-application with 1 µM DS2 (gray trace). **B.** γ2 presence was validated by 1 µM Zn^2+^ sensitivity (light blue trace). YX84 (5 µM) were applied in both Panel A and B (red trace). **C.** Peak YX84 current was compared to GABA current. α4/δ subunit-containing receptors exhibited greater efficacy relative to GABA compared to α1/ γ2 subunit-containing receptors (p=0.0034).

### 3.2 Pharmacological Profile of YX84-Gated Currents

Pharmacological dissection of YX84-gated currents revealed partial sensitivity to bicuculline and weak sensitivity to gabazine, suggesting a non-orthosteric binding site. (Figure 6A-B) (Ueno et al., 1997). The combination of YX84 potentiation, gating, and block evokes similarities with barbiturate actions (Evans, 1979; Akk and Steinbach, 2000).

**Figure 6.**
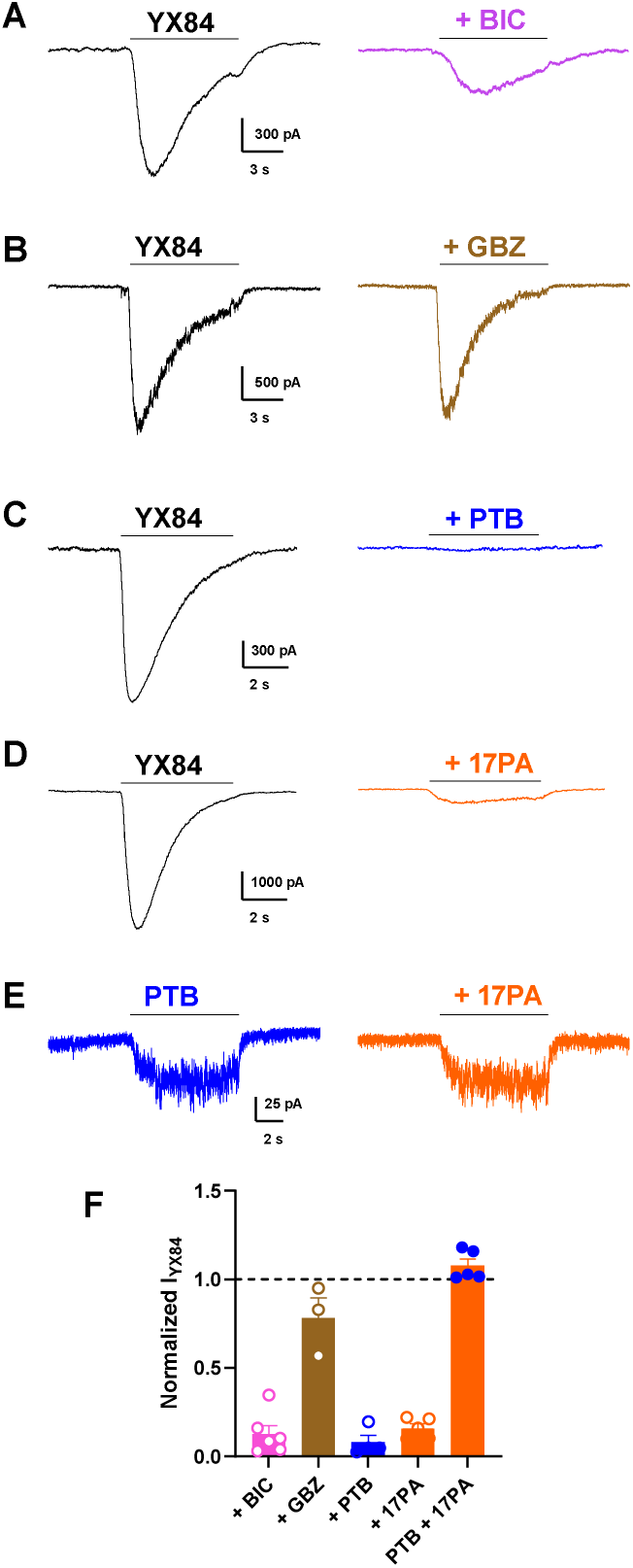
Effect of GABA_A_ ligands on YX84-gated current. **A.** YX84 (5 µM) current is partially sensitive to bicuculline (100 µM). **B.** YX84 is very weakly sensitive to gabazine (10 µM). Pentobarbital (PTB, 50 µM occludes the YX84 current). The neurosteroid antagonist 17PA (10 µM) inhibits the current gated by YX84 (**D**) but not that gated by PTB (**E**). **F.** Summary of experiments shown in A-E.

Consistent with this, pentobarbital occluded YX84 responses (Figure 6C). On the other hand, the neurosteroid antagonist 17PA inhibited YX84-gated currents but not those gated by pentobarbital (Figure 6D, E), supporting a distinct neurosteroid-sensitive site of action.

### 3.3. YX84 Exhibits Weak NMDA Receptor Inhibition

YX84 modestly inhibited NMDA-evoked currents in hippocampal neurons, with an estimated IC₅₀ in the low micromolar range (Figure 7). The inhibition was incomplete, suggesting weak negative allosteric modulation rather than channel blockade.

**Figure 7.**
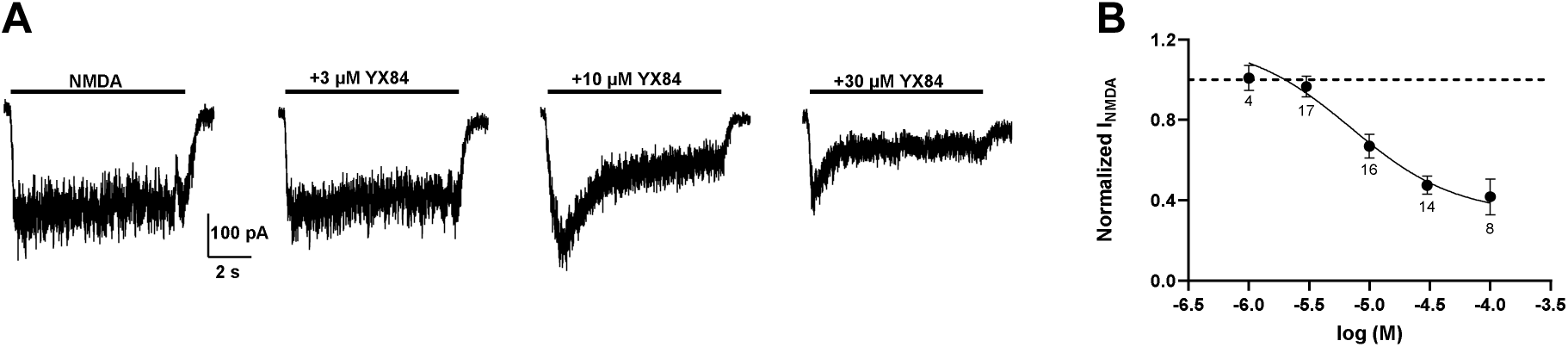
YX84 is a weak NMDAR inhibitor. **A.** Responses of a hippocampal neuron to NMDA (10 µM) alone (left) and co-applied with increasing YX84 concentrations. **B.** Summary of the effect of YX84 relative to response gated by NMDA alone. Based on a fit to averaged data, the IC_50_ was 7.02 µM and predicted maximum effect was 0.33.

### 3.4 YX84 Does Not Impair Sensorimotor Function

As an initial assay to assess potential behavioral effects, we tested YX84 (30 mg/kg, i.p.) in a battery of sensorimotor assays. Unlike allopregnanolone, which impaired balance and strength measures, YX84 had no detectable effects on any behavioral endpoint (Figure 8). The results suggest a challenge of brain penetration and/or rapid metabolism, but the results also indicate an encouraging safety profile.

**Figure 8.**
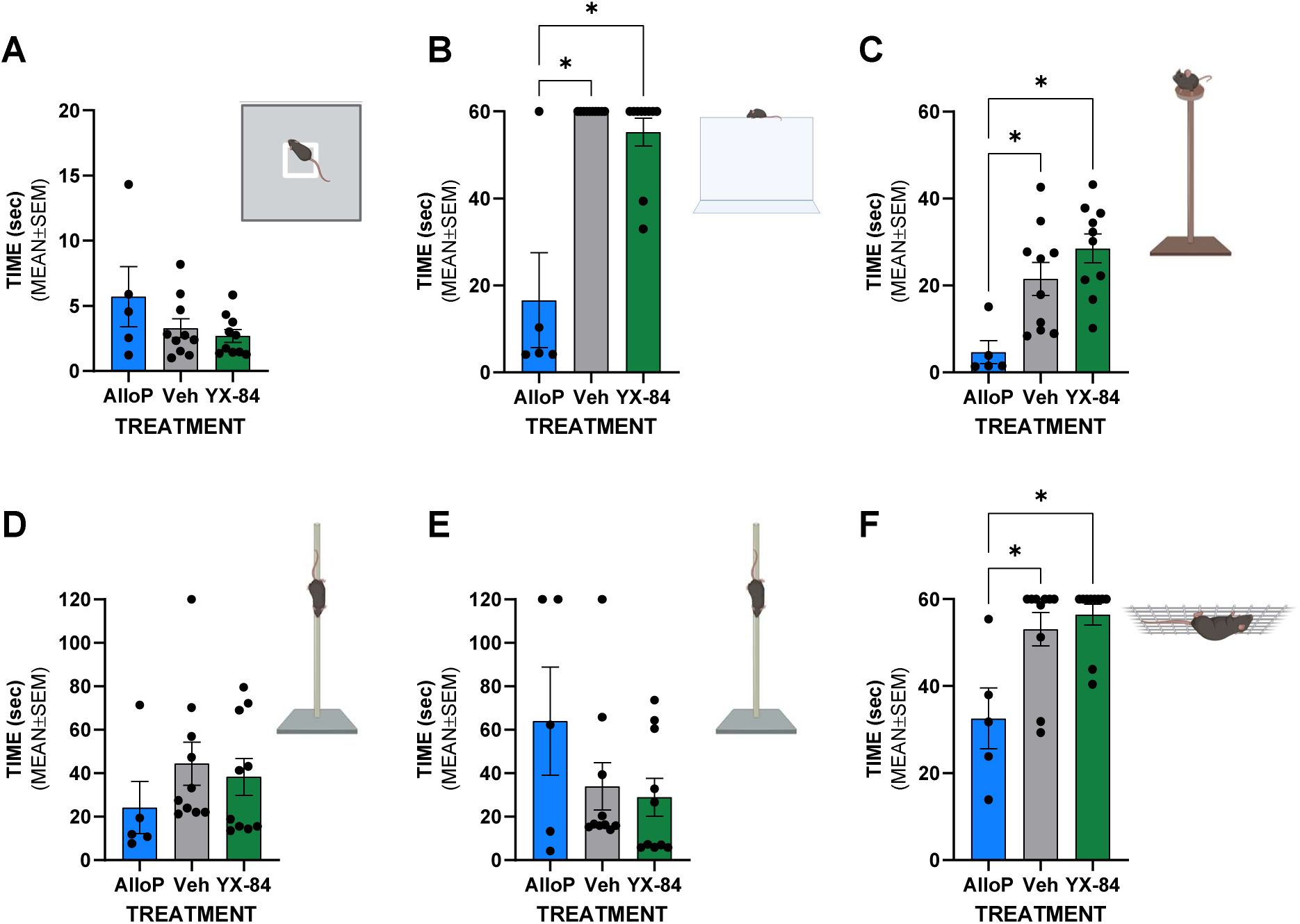
Lack of acute sensorimotor effects of ip YX84 (30 mg/kg). Allopregnanolone (AlloP; 15 mg/kg) was used as a positive control. As indicated by the drawings, the battery included measures of walk initiation (**A**), ledge walking and balancing (**B**), platform balancing (**C**), pole climbing fine motor test (**D**), pole reversal (**E**), and inverted screen strength (**F**). Allopregnanolone significantly reduced balance challenges and inverted screen behaviors. Meanwhile, YX84 had no effect on any of the measures.

## 4. Discussion

Our findings reveal that YX84 and related analogues represent a novel class of neuroactive steroids with non-classical structure-activity relationships and an unusual profile for potentiation and gating. Although YX84 adheres to a typical pattern in which low concentrations exhibit positive allosteric modulation and higher concentrations gate the receptor (Figure 1), the efficacy for gating greatly exceeded that of canonical neurosteroids and approached or exceeded that of the native neurotransmitter. The requirement for both the sulfate and trifluoromethyl ketone groups, along with the lack of stereoselectivity and C5 reduction, challenges existing models of neurosteroid action at GABA_A_ receptors. These results suggest that YX84 engages a distinct modulatory site or mechanism, potentially offering new avenues for therapeutic targeting.

Another hint of a site distinct from existing neurosteroids is given by the rapid onset of gating (Figure 1) relative to allopregnanolone (Shu et al., 2004). The slow onset of allopregnanolone gating has been attributed to slow membrane partitioning. Rapid gating by YX84 and all analogues depicted in Figure 2A appears more akin to binding governed by aqueous diffusion and may suggest direct access to receptor binding sites. Aqueous access with limited membrane partitioning may be predicted from the charged sulfated group associated with active analogues (Figure 2A).

Efficacy of YX84 was comparable or greater than that of GABA on native receptors in mouse hippocampal neurons, recombinant α1β2γ2 and α4β2δ receptors (Figures 3, 5). The efficacy at δ-containing receptors may be especially surprising, achieving more than 5-fold greater response than a saturating GABA concentration in some transfected N2a cells (Figure 5C). The results are reminiscent of those achieved by the ‘super agonist’ THIP, which has seen therapeutic interest for insomnia and neurodevelopment disorders (Wafford and Ebert, 2006; Samanta et al., 2025). THIP exhibits ∼2-fold greater efficacy compared with GABA in past studies (Mortensen et al., 2010). Because agonists cannot gate channels to an open probability greater than 1, our results showing 5-fold potentiation (Figure 5C) would suggest that GABA efficacy at α4β2δ receptors expressed in N2a cells is very low, with a peak channel open probability ∼0.2 (Figure 5C). Strong DS2 potentiation of GABA responses is also consistent with a low basal open probability at 10 µM GABA (Figure 5A).

An alternative or additional explanation for the apparent large YX84 efficacy value could be an increase in single-channel conductance level.

Gating by YX84 is followed by apparent desensitization, which is faster and more profound than exhibited by the native agonist GABA. This difference is reminiscent of classical agonist dependent differences in desensitization properties of different agonists at AMPA-type glutamate receptors (Patneau and Mayer, 1991; Thio et al., 1991). However, unlike the glutamate receptor analogy, YX84 and GABA do not act at a common orthosteric site, based on antagonist pharmacology in Figure 6. Results in Figure 4 are consistent with a channel blocking role for YX84 in the apparent desensitization phenomenon.

Results of antagonist challenge and occlusion (Figure 6) suggest that YX84 exhibits a unique pattern of binding and receptor activation. The pattern of strong but incomplete bicuculline inhibition and weak gabazine inhibition is reminiscent of other allosteric gating profiles, including those of barbiturates and neurosteroids (Ueno et al., 1997). However, the combination of 17PA and pentobarbital inhibition of YX84 gating was unexpected because 17PA has been proposed to be a selective inhibitor of neurosteroid potentiation (Mennerick et al., 2004; Kelley et al., 2007). Taken together, the results suggest that YX84 overlaps both neurosteroid and pentobarbital binding sites. Alternative explanations for the 17PA effect on YX84 gating could be sequestration in solution of active compound through micelle formation, competitive binding to solubilizers (DMSO in our case), or steroid-steroid stacking between YX84 and 17PA in solution. However, these possibilities enjoy limited support from the literature at concentrations employed.

Although our initial behavioral screen did not reveal detectable sensorimotor impairment following YX84 administration, these findings provide important context for interpreting the compound’s pharmacological profile. Despite exhibiting strong GABA_A_ receptor agonism in vitro, including high efficacy at both synaptic and extrasynaptic receptor populations, YX84 failed to elicit the motor slowing, balance deficits, or strength impairments commonly associated with neuroactive steroid–induced CNS depression. In contrast, allopregnanolone produced clear impairments across multiple tasks in the same battery, highlighting the relative behavioral sparing of YX84 at doses that are pharmacologically active in electrophysiological assays. This divergence suggests that brain exposure to YX84 may be limited by poor permeability or rapid metabolism, consistent with predictions based on its sulfated structure, but it also indicates a lack of prominent off-target peripheral adverse effects affecting measured behaviors. The absence of acute sensorimotor disruption thus reduces concerns regarding peripheral toxicity and supports the feasibility of further exploration into delivery strategies and CNS-targeted applications of this structurally non-classical neuroactive steroid.

The dual activity, potent GABA_A_ receptor activation and weak NMDA receptor inhibition, is not unique (Mennerick et al., 2001; Ziolkowski et al., 2021; Abramova et al., 2023) but may be particularly advantageous in disorders characterized by excitatory/inhibitory imbalance, such as epilepsy or mood disorders. The lack of acute sensorimotor impairment at behaviorally relevant doses could support the feasibility of further development and a lack of off-target peripheral effects. On the other hand, these results are likely driven by the limited blood-brain permeability expected from charged neurosteroids. Effective CNS treatment strategies will likely require consideration of effective delivery to brain and further tests of side effects.

Future studies will explore subunit selectivity, pharmacokinetics, and therapeutic efficacy in disease models. These efforts may clarify the translational potential of YX84 and its analogues as multifunctional modulators of synaptic transmission.

## Supporting information

Supplemental chemistry methods

## Author contributions

Conceptualization: Douglas F. Covey, Steven Mennerick, Charles F. Zorumski, Hong-Jin Shu. Metholology: Yuanjian Xu, Hong-Jin Shu, Mingxing Qian, Carla M. Yuede, Ann Benz. Formal Analysis: Douglas F. Covey, Hong-Jin Shu, Carla M. Yuede, Steven Mennerick. Writing, original draft: Steven Mennerick. Writing, review and editing: all authors. Funding acquisition: Douglas F. Covey, Steven Mennerick, Charles F. Zorumski

## Acknowledgements

The authors thank members of the Taylor Family Institute for Innovative Psychiatric Research for discussion, particularly Dr. Alex Evers for critical feedback on the manuscript. This work was supported in part by R01 MH123748 (SM) and P50 MH122379 (SM and CFZ). The Animal Behavior Core is supported in part by funds from the McDonnell Center for Systems Neuroscience and the Taylor Family Institute. During the preparation of this work, the authors used generative artificial intelligence tools to aid grammar and stylistic flow. The authors reviewed and edited the content and take full responsibility for the content of the publication.

## Data availability statement

Data supporting the findings of this study are available from the authors upon reasonable request.

