## Supplemental chemistry methods for "A Non-Classical Neuroactive Steroid Exhibiting Potent, Efficacious GABA_A_ Receptor Agonism and NMDA Receptor Inhibition"

### Synthesis of YX77 and YX78

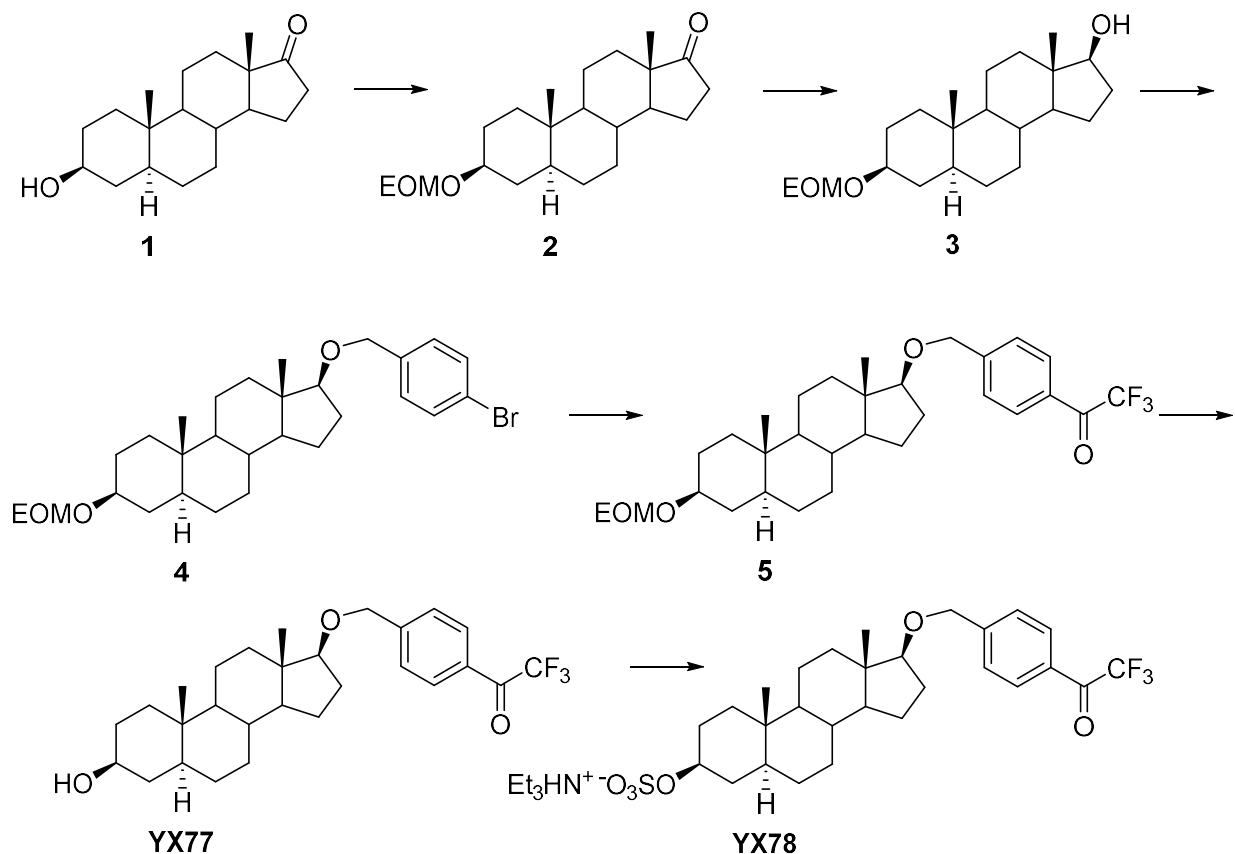

**(3β,5α)-3-(Ethoxymethoxy)-androst-17-one (2).** To a stirred solution of epiandrosterone (**1**, 580 mg, 2 mmol) in $\text{CH}_2\text{Cl}_2$ (30 mL) was added chloromethyl ethyl ether (0.4 mL, 4 mmol) and $(i\text{-Pr})_2\text{NEt}$ (0.8 mL, 6 mmol) at 23 °C. The reaction was stirred for 16 h. $\text{CH}_2\text{Cl}_2$ was removed and the residue purified by flash column chromatography (silica gel, eluted with 10% EtOAc in hexanes) to give steroid **2** (650 mg, 94%): $^1\text{H}$ NMR (400 MHz, $\text{CDCl}_3$) $\delta$ 4.68 (q, $J$ = 6.6 Hz, 2H), 3.58 (q, $J$ = 7.0 Hz, 2H), 3.49-3.44 (m, 1H), 2.30 (dd, $J$ = 8.6 Hz, 10.5 Hz, 1H), 2.06-0.61 (m, 21H), 1.18 (t, $J$ = 7.0 Hz, 3H), 0.81 (s, 3H), 0.78 (s, 3H); $^{13}\text{C}$ NMR (100 MHz, $\text{CDCl}_3$) $\delta$

221.2, 93.0, 75.9, 62.9, 54.4, 51.3, 47.7, 44.8, 36.9, 35.8, 35.7, 35.1, 35.0, 31.5, 30.9, 28.6, 28.4, 21.7, 20.4, 15.1, 13.8, 12.2.

**(3β,5α,17β)-3-(Ethoxymethoxy)-androstan-17-ol (3).** To a stirred solution of steroid **2** (650 mg, 1.86 mmol) in EtOH (20 mL) was added NaBH<sub>4</sub> (200 mg, 5.3 mmol) at 23 °C. After 1 h, aqueous NaHCO<sub>3</sub> was added and stirring continued for 15 min. Most of ethanol was removed, water (50 mL) was added to the remaining aqueous solution and the product was extracted into EtOAc (100 mL x 2). The combined extracts were washed with brine (100 mL x 3), water (100 mL), dried over anhydrous Na<sub>2</sub>SO<sub>4</sub>, filtered and the solvent was removed to give steroid **3** (630 mg, 97%): <sup>1</sup>H NMR (400 MHz, CDCl<sub>3</sub>) $\delta$ 4.72 (q, $J$ = 6.6 Hz, 2H), 3.61-3.56 (m, 3H), 3.53-3.45 (m, 1H), 2.06-1.97 (m, 1H), 1.83-0.57 (m, 22H), 1.21 (t, $J$ = 7.0 Hz, 3H), 0.79 (s, 3H), 0.71 (s, 3H); <sup>13</sup>C NMR (100 MHz, CDCl<sub>3</sub>) $\delta$ 93.0, 81.8, 76.0, 83.0, 54.5, 51.0, 44.9, 43.0, 37.0, 36.7, 35.7, 35.5, 35.2, 31.6, 30.4, 28.6 (2 x C), 23.4, 20.8, 15.1, 12.3, 11.1.

**(3β,5α,17β)-17-((4-Bromobenzyl)oxy)-3-(ethoxymethoxy)-androsterane (4).** To a stirred solution of steroid **3** (630 mg, 1.79 mmol) in THF (30 mL) was added KH (600 mg, 30% in mineral oil, 4.5 mmol). The mixture was refluxed for 30 min. 4-Bromobenzyl bromide (1.0 g, 4 mmol) in THF (10 mL) was added and the reaction was refluxed for 16 h. After cooling, water was added and the product was extracted into EtOAc (100 mL x 2). The combined extracts were dried over anhydrous Na<sub>2</sub>SO<sub>4</sub>, filtered, the solvent removed and the residue purified by flash column chromatography (silica gel, eluted with 2-5% EtOAc in hexanes) to give steroid **4** (800 mg, 86%): <sup>1</sup>H NMR (400 MHz, CDCl<sub>3</sub>) $\delta$ 7.45 (d, $J$ = 8.1 Hz, 2H), 7.20 (d, $J$ = 8.1 Hz, 2H), 4.72

(q, $J = 7.0$ Hz, 2H), 4.46 (s, 2H), 3.62 (q, $J = 7.0$ Hz, 2H), 3.60 (s, 1H), 3.38-3.33 (m, 1H), 1.99-0.57 (m, 22H), 1.22 (t, $J = 7.0$ Hz, 3H), 0.80 (s, 3H), 0.80 (s, 3H); $^{13}\text{C}$ NMR (100 MHz, $\text{CDCl}_3$) $\delta$ 138.4, 131.3 (2 x C), 128.9 (2 x C), 121.0, 93.0, 88.5, 76.0, 70.8, 63.0, 54.5, 51.2, 44.9, 43.1, 38.0, 37.0, 35.7, 35.3, 35.2, 31.6, 28.6 (2 x C), 27.9, 23.4, 20.9, 15.2, 12.3, 11.9.

**1-(4-(((3 $\beta$ -(Ethoxymethoxy)-5 $\alpha$ -androstan-17 $\beta$ -yl)oxy)methyl)phenyl)-2,2,2-trifluoroethan-**

**1-one (5).** To a stirred solution of steroid **4** (800 mg, 1.54 mmol) in THF (30 mL) was slowly added *n*-BuLi (2.5 M in THF, 2 mL, 5 mmol) at $-78^\circ\text{C}$. After 45 min, 1-trifluroacetyl piperidine (1.3 g, 7 mmol) was added and the mixture was stirred at $-78^\circ\text{C}$ for 1 h. Aqueous $\text{NaHCO}_3$ (10 mL) was added at $-78^\circ\text{C}$ and the reaction slowly warmed to $23^\circ\text{C}$. The product was extracted into EtOAc (2 x 100 mL). The combined extracts were dried over anhydrous $\text{Na}_2\text{SO}_4$, filtered, the solvent removed and the residue purified by flash column chromatography (silica gel, eluted with 5-10% EtOAc in hexanes) to give steroid **5** (470 mg, 57%): $^1\text{H}$ NMR (400 MHz, $\text{CDCl}_3$) $\delta$ 8.04 (d, $J = 8.2$ Hz, 2H), 7.51 (d, $J = 8.2$ Hz, 2H), 4.74 (q, $J = 7.0$ Hz, 2H), 4.60 (s, 2H), 3.62 (q, $J = 7.0$ Hz, 2H), 3.57-3.47 (m, 1H), 3.42 (t, $J = 8.2$ Hz, 1H), 2.03-0.58 (m, 22H), 1.22 (t, $J = 7.0$ Hz, 3H), 0.82 (s, 3H), 0.81 (s, 3H); $^{13}\text{C}$ NMR (100 MHz, $\text{CDCl}_3$) $\delta$ 180.6 (q, $J = 35.1$ Hz), 148.1, 130.1, 130.1, 128.7, 127.2 (2 x C), 121.0 (q, $J = 291.4$ Hz), 93.0, 89.0, 76.0, 70.6, 62.9, 54.4, 51.1, 44.8, 43.2, 37.9, 37.0, 35.6, 35.2, 35.2, 31.6, 28.6, 28.6, 27.8, 23.3, 20.8, 15.1, 12.2, 11.8.

**1-(4-(((3 $\beta$ -Hydroxy-5 $\alpha$ -androstan-17 $\beta$ -yl)oxy)methyl)phenyl)-2,2,2-trifluoroethan-1-one**

**(YX77).** To steroid **5** (470 mg, 0.88 mmol) was added THF (10 mL) and 6 N HCl (10 mL) at $23^\circ\text{C}$. After stirring for 2 h, water was added and the product was extracted into $\text{CH}_2\text{Cl}_2$ (2 x 150

mL). The combined extracts were washed with aqueous NaHCO<sub>3</sub>, dried over anhydrous Na<sub>2</sub>SO<sub>4</sub>, filtered, the solvent removed and the residue purified by flash column chromatography (silica gel, eluted with 20% EtOAc in hexanes) to give **YX77** (410 mg, 98%): <sup>1</sup>H NMR (400 MHz, CDCl<sub>3</sub>) δ 8.04 (d, *J* = 8.1 Hz, 2H), 7.50 (d, *J* = 8.1 Hz, 2H), 4.61 (s, 2H), 3.58-3.53 (m, 1H), 3.42 (t, *J* = 8.1 Hz, 1H), 2.03-0.57 (m, 23H), 0.82 (s, 3H), 0.80 (s, 3H); <sup>13</sup>C NMR (100 MHz, CDCl<sub>3</sub>) δ 180.3 (q, *J* = 35.1 Hz), 148.1, 130.2, 130.2, 128.7, 127.2 (2 x C), 121.0 (q, *J* = 291.5 Hz), 89.1, 71.2, 70.6, 54.5, 51.2, 44.9, 43.2, 38.1, 38.0, 37.0, 35.5, 35.3, 31.6, 31.4, 28.5, 27.8, 23.4, 20.9, 12.3, 11.9.

**1-(4-(((3β-Sulfoxy-5α-androstan-17β-yl)oxy)methyl)phenyl)-2,2,2-trifluoroethan-1-one, triethyl ammonium salt (YX78).** To a stirred solution of **YX77** (60 mg, 0.13 mmol) in pyridine (2 mL) was added SO<sub>3</sub>-Me<sub>3</sub>N (50 mg, 0.36 mmol) at 23 °C. The mixture was stirred at 23 °C for 20 h. Pyridine was removed under reduced pressure and the residue was purified by flash chromatography (silica gel, eluted with CH<sub>2</sub>Cl<sub>2</sub>/MeOH/Et<sub>3</sub>N = 100/10/1 (v/v/v)) to give **YX78** (43 mg, 50%): <sup>1</sup>H NMR (400 MHz, CDCl<sub>3</sub>) δ 8.03 (d, *J* = 7.4 Hz, 2H), 7.50 (d, *J* = 7.8 Hz, 2H), 4.60 (s, 2H), 4.33-4.28 (m, 1H), 3.41 (t, *J* = 7.6 Hz, 1H), 3.16 (q, *J* = 6.6 Hz, 6H), 2.02-0.60 (m, 23H), 1.36 (t, *J* = 7.0 Hz, 9H), 0.80 (s, 3H), 0.79 (s, 3H); <sup>13</sup>C NMR (100 MHz, CDCl<sub>3</sub>) δ 180.3 (q, *J* = 35.1 Hz), 148.1, 130.2, 130.2, 128.7, 127.2 (2 x C), 118.1 (q, *J* = 311.5 Hz), 89.1, 78.2, 70.6, 54.3, 51.1, 46.4 (3 x C), 44.9, 43.2, 37.9, 37.0, 35.4, 35.2, 35.1, 31.6, 28.6, 28.4, 27.8, 23.4, 20.8, 12.2, 11.9, 8.7 (3 x C).

### Synthesis of YX79 and YX80

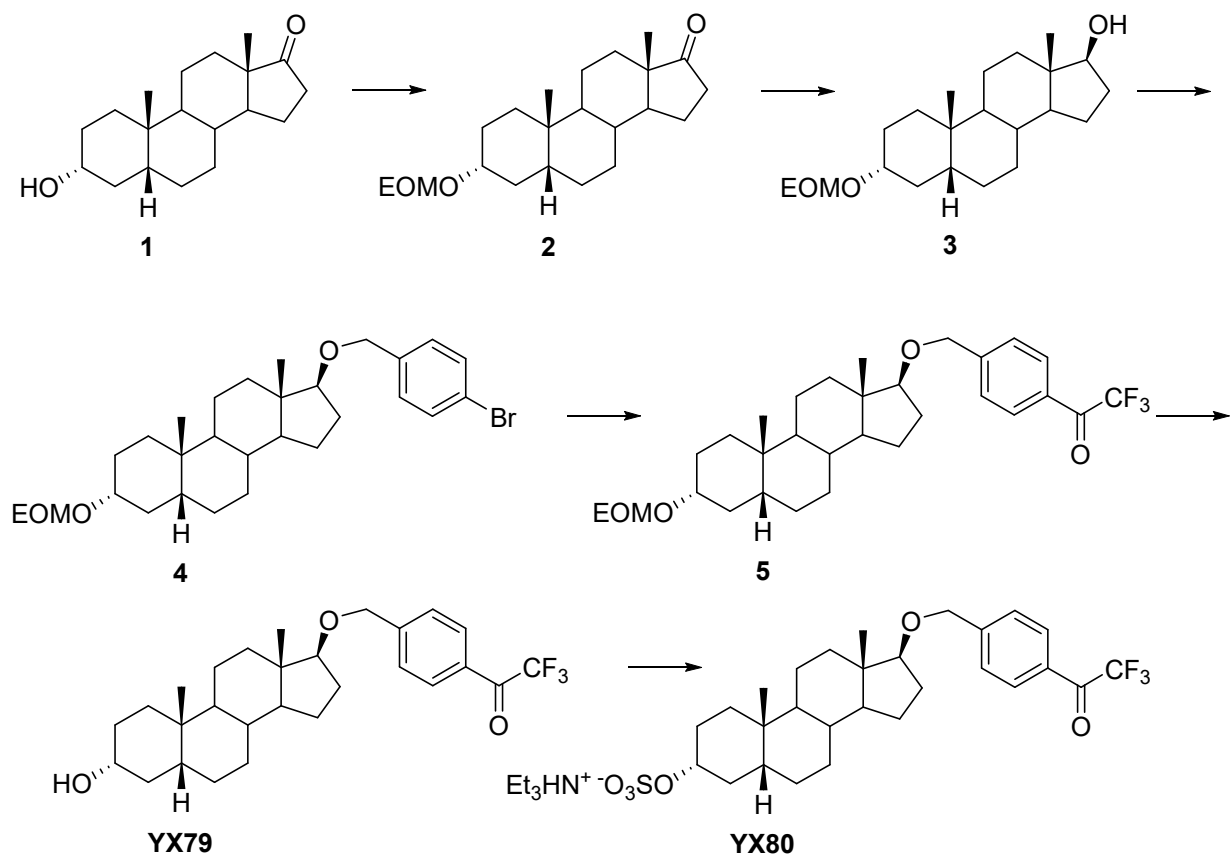

**(3α,5β)-3-(Ethoxymethoxy)-androst-17-one (2).** To a stirred solution of etiocholanolone (**1**, 580 mg, 2 mmol) in $\text{CH}_2\text{Cl}_2$ (30 mL) was added chloromethyl ethyl ether (0.4 mL, 4 mmol) and $(i\text{-Pr})_2\text{NEt}$ (0.8 mL, 6 mmol) at 23 °C. The reaction was stirred at 23 °C for 16 h. $\text{CH}_2\text{Cl}_2$ was removed and the residue purified by column flash chromatography (silica gel, eluted with 10% EtOAc in hexanes) to give steroid **2** (640 mg, 92%): $^1\text{H}$ NMR (400 MHz, $\text{CDCl}_3$) $\delta$ 4.73 (q, $J$ = 6.6 Hz, 2H), 3.64–3.53 (m, 3H), 2.47 (dd, $J$ = 8.6 Hz, 10.5 Hz, 1H), 2.13–0.99 (m, 21H), 1.23 (t, $J$ = 7.0 Hz, 3H), 0.95 (s, 3H), 0.85 (s, 3H); $^{13}\text{C}$ NMR (100 MHz, $\text{CDCl}_3$) $\delta$ 221.4, 93.1, 76.5,

63.0, 51.5, 47.9, 42.0, 40.7, 35.9, 35.4, 35.3, 34.9, 33.5, 31.7, 27.6, 26.9, 25.3, 23.3, 21.8, 20.1, 15.2, 13.8.

**(3α,5β,17β)-3-(Ethoxymethoxy)-androstan-17-ol (3).** To a stirred solution of steroid **2** (640 mg, 1.83 mmol) in EtOH (20 mL) was added NaBH<sub>4</sub> (200 mg, 5.3 mmol) at 23 °C. After 1 h, aqueous NaHCO<sub>3</sub> was added and the reaction was stirred for 15 min. Most of ethanol was removed and to the remaining aqueous solution water (50 mL) was added. The product was extracted into EtOAc (100 mL x 2). The combined extracts were washed with brine (100 mL x 3), water (100 mL), dried over anhydrous Na<sub>2</sub>SO<sub>4</sub>, filtered and the solvent removed to give steroid **3** (630 mg, 98%): <sup>1</sup>H NMR (400 MHz, CDCl<sub>3</sub>) $\delta$ 4.72 (q, $J$ = 6.6 Hz, 2H), 3.64-3.50 (m, 4H), 2.06-1.99 (m, 1H), 1.87-0.87 (m, 22H), 1.22 (t, $J$ = 7.0 Hz, 3H), 0.92 (s, 3H), 0.71 (s, 3H); <sup>13</sup>C NMR (100 MHz, CDCl<sub>3</sub>) $\delta$ 93.1, 81.9, 76.6, 63.0, 51.1, 43.1, 42.1, 40.6, 36.9, 35.9, 35.4, 34.8, 33.5, 30.6, 27.6, 27.1, 26.0, 23.4 (2 x C), 20.4, 15.2, 11.1.

**(3α,5β,17β)-17-((4-Bromobenzyl)oxy)-3-(ethoxymethoxy)-androstande (4).** To a stirred solution of steroid **3** (630 mg, 1.79 mmol) in THF (30 mL) was added KH (600 mg, 30% in mineral oil, 4.5 mmol). The mixture was refluxed for 30 min. 4-Bromobenzyl bromide (1.0 g, 4 mmol) in THF (10 mL) was added and the reaction was refluxed for 16 h. After cooling, water was added and the product was extracted into EtOAc (100 mL x 2), dried over anhydrous Na<sub>2</sub>SO<sub>4</sub>, filtered, the solvent removed and the residue purified by flash column chromatography (silica gel, eluted with 2-5% EtOAc in hexanes) to give steroid **4** (870 mg, 94%): <sup>1</sup>H NMR (400 MHz, CDCl<sub>3</sub>) $\delta$ 7.45 (d, $J$ = 8.2 Hz, 2H), 7.21 (d, $J$ = 8.1 Hz, 2H), 4.73 (q, $J$ = 7.0 Hz, 2H), 4.47

(s, 2H), 3.63-3.51 (m, 3H), 3.40 (t, $J = 8.2$ Hz, 1H), 2.04-0.95 (m, 22H), 1.23 (t, $J = 7.0$ Hz, 3H), 0.92 (s, 3H), 0.79 (s, 3H); $^{13}\text{C}$ NMR (100 MHz, $\text{CDCl}_3$) $\delta$ 138.4, 131.3 (2 x C), 128.9 (2 x C), 121.0, 93.0, 88.7, 76.6, 70.9, 63.0, 51.2, 43.2, 42.1, 40.6, 38.1, 35.6, 35.4, 34.8, 33.5, 28.0, 27.7, 27.1, 25.9, 23.4 (2 x C), 20.4, 15.2, 11.8.

**1-(4-(((3α-(Ethoxymethoxy)-5β-androst-17β-yl)oxy)methyl)phenyl)-2,2,2-trifluoroethan-1-one (5).** To a stirred solution of steroid **4** (870 mg, 1.68 mmol) in THF (30 mL) was slowly added *n*-BuLi (2.5 M in THF, 2 mL, 5 mmol) at $-78$ °C. After 45 min, 1-trifluroacetyl piperidine (1.3 g, 7 mmol) was added and the reaction was stirred at $-78$ °C for 1 h. Aqueous $\text{NaHCO}_3$ (10 mL) was added at $-78$ °C and the reaction was slowly warmed to $23$ °C. The product was extracted into EtOAc (2 x 100 mL). The combined extracts were dried over anhydrous $\text{Na}_2\text{SO}_4$, filtered, the solvent removed and the residue purified by flash column chromatography (silica gel, eluted with 5-10% EtOAc in hexanes) to give steroid **5** (570 mg, 63%): $^1\text{H}$ NMR (400 MHz, $\text{CDCl}_3$) $\delta$ 8.05 (d, $J = 7.8$ Hz, 2H), 7.51 (d, $J = 8.2$ Hz, 2H), 4.72 (q, $J = 7.0$ Hz, 2H), 4.61 (s, 2H), 3.62-3.51 (m, 3H), 3.43 (t, $J = 8.2$ Hz, 1H), 2.04-0.96 (m, 22H), 1.22 (t, $J = 7.0$ Hz, 3H), 0.92 (s, 3H), 0.81 (s, 3H); $^{13}\text{C}$ NMR (100 MHz, $\text{CDCl}_3$) $\delta$ 180.3 (q, $J = 34.4$ Hz), 148.1, 130.21, 130.19, 128.7, 127.2 (2 x C), 118.2 (q, $J = 291.5$ Hz), 93.1, 89.2, 76.6, 70.7, 63.0, 51.2, 43.3, 42.1, 40.6, 38.1, 35.6, 35.4, 34.8, 33.5, 27.9, 27.7, 27.1, 25.9, 23.42, 23.38, 20.4, 15.2, 11.8.

**1-(4-(((3α-Hydroxy-5β-androst-17β-yl)oxy)methyl)phenyl)-2,2,2-trifluoroethan-1-one (YX79).** To steroid **5** (570 mg, 1.06 mmol) was added THF (10 mL) and 6 N HCl (10 mL) at $23$ °C. After stirring for 2 h, water was added and the product was extracted into $\text{CH}_2\text{Cl}_2$ (2 x 150

mL). The combined extracts were washed with aqueous NaHCO<sub>3</sub>, dried over anhydrous Na<sub>2</sub>SO<sub>4</sub>, filtered, the solvent removed and the residue purified by flash column chromatography (silica gel, eluted with 20% EtOAc in hexanes) to give **YX79** (470 mg, 93%): <sup>1</sup>H NMR (400 MHz, CDCl<sub>3</sub>) δ 8.03 (d, *J* = 7.8 Hz, 2H), 7.50 (d, *J* = 8.2 Hz, 2H), 4.60 (s, 2H), 3.62-3.57 (m, 1H), 3.42 (t, *J* = 8.1 Hz, 1H), 2.13-0.85 (m, 23H), 0.90 (s, 3H), 0.80 (s, 3H); <sup>13</sup>C NMR (100 MHz, CDCl<sub>3</sub>) δ 180.6 (q, *J* = 35.1 Hz), 148.1, 130.19, 130.17, 128.7, 127.2 (2 x C), 121.0 (q, *J* = 291.5 Hz), 89.1, 71.6, 70.6, 51.2, 43.3, 42.0, 40.6, 38.1, 36.3, 35.6, 35.4, 34.6, 30.4, 27.9, 27.0, 26.0, 23.4, 23.3, 20.4, 11.8.

**1-(4-(((3 $\alpha$ -Sulfoxy-5 $\beta$ -androst-17 $\beta$ -yl)oxy)methyl)phenyl)-2,2,2-trifluoroethan-1-one,**

**triethyl ammonium salt (YX80).** To a stirred solution of **YX79** (163 mg, 0.34 mmol) in pyridine (5 mL) was added SO<sub>3</sub>-Me<sub>3</sub>N (120 mg, 0.86 mmol) at 23 °C. The mixture was stirred at 23 °C for 20 h. Pyridine was removed under reduced pressure and the residue was purified by flash column chromatography (silica gel, eluted with CH<sub>2</sub>Cl<sub>2</sub>/MeOH/Et<sub>3</sub>N = 100/10/1 (v/v/v)) to give **YX80** (184 mg, 79%): <sup>1</sup>H NMR (400 MHz, CDCl<sub>3</sub>) δ 7.99 (d, *J* = 8.2 Hz, 2H), 7.47 (d, *J* = 8.2 Hz, 2H), 4.57 (s, 2H), 4.32-4.27 (m, 1H), 3.39 (t, *J* = 7.8 Hz, 1H), 3.15 (q, *J* = 7.4 Hz, 6H), 2.00-0.79 (m, 23H), 1.34 (t, *J* = 7.4 Hz, 9H), 0.87 (s, 3H), 0.76 (s, 3H); <sup>13</sup>C NMR (100 MHz, CDCl<sub>3</sub>) δ 180.7 (q, *J* = 35.1 Hz), 148.2, 130.2, 130.1, 128.6, 127.2 (2 x C), 121.0 (q, *J* = 291.4 Hz), 89.1, 74.7, 70.6, 51.2, 46.4 (3 x C), 43.3, 42.1, 40.5, 38.1, 35.5, 35.3, 37.5, 33.3, 27.8, 27.7, 26.8, 25.9, 23.4, 23.3, 20.4, 11.8, 8.7 (3 x C).

Chemical reaction scheme showing the synthesis of YX81 and YX82 from compound 1.

The scheme consists of two rows of reactions:

Row 1: Compound 1 (a steroid with a ketone at C3 and a hydroxyl group at C14) is converted to compound 2 (a steroid with a ketone at C3 and a hydroxyl group at C14, with a methyl group at C13). Compound 2 is converted to compound 3 (a steroid with a ketone at C3 and a hydroxyl group at C14, with a methyl group at C13). Compound 3 is converted to compound 4 (a steroid with a ketone at C3 and a hydroxyl group at C14, with a methyl group at C13).

Row 2: Compound 4 is converted to compound 5 (a steroid with a ketone at C3 and a hydroxyl group at C14, with a methyl group at C13). Compound 5 is converted to compound 6 (a steroid with a ketone at C3 and a hydroxyl group at C14, with a methyl group at C13). Compound 6 is converted to compound 7 (a steroid with a ketone at C3 and a hydroxyl group at C14, with a methyl group at C13). Compound 7 is converted to compound 8 (a steroid with a ketone at C3 and a hydroxyl group at C14, with a methyl group at C13). Compound 8 is converted to YX81 (a steroid with a ketone at C3 and a hydroxyl group at C14, with a methyl group at C13). YX81 is converted to YX82 (a steroid with a ketone at C3 and a hydroxyl group at C14, with a methyl group at C13).

**(5β,17β)-17-Hydroxyandrostane-3-one (2).** To a solution of testosterone (**1**, 2 g, 6.94 mmol) in pyridine (20mL) was added Pd/C (10%, 200 mg) in a Parr hydrogenation flask and hydrogenation was carried out at 55 psi for 4 h. The mixture was filtered through celite and washed with EtOAc. The solvent was removed and the residue was purified by flash column chromatography (silica gel, eluted with 20 % EtOAc in hexanes) to give steroid **2** (1.8 g, 90%):

$^1\text{H}$ NMR (400 MHz, $\text{CDCl}_3$) $\delta$ 3.67-3.63 (m, 1H), 2.70-2.63 (m, 1H), 2.36-2.27 (m, 1H), 2.17-1.05 (m, 21H), 1.02 (s, 3H), 0.75 (s, 3H); $^{13}\text{C}$ NMR (100 MHz, $\text{CDCl}_3$) $\delta$ 213.3, 81.8, 61.0, 44.3, 43.1, 42.3, 40.9, 37.2, 37.0, 36.8, 35.6, 35.0, 30.5, 26.5, 25.4, 23.4, 22.7, 20.8, 11.1.

**5β-Androstane-3,17-dione (3).** To a stirred solution of steroid **2** (1.8 g, 6.2 mmol) in acetone (20 mL) was added Jones's reagent at 0 °C until a yellowish brown color persisted. After 10 min, 2-propanol (2 mL) and then water (50 mL) were added. The product was extracted into EtOAc (150 mL x 3). The combined extracts were dried over anhydrous $\text{Na}_2\text{SO}_4$, filtered, the solvent removed and the residue purified by flash column chromatography (silica gel, eluted with 20 % EtOAc in hexanes) to give steroid **3** (1.65 g, 92%): $^1\text{H}$ NMR (400 MHz, $\text{CDCl}_3$) $\delta$ 2.68-2.61 (m, 1H), 2.47-2.40 (m, 1H), 2.33-1.16 (m, 20H), 1.02 (s, 3H), 0.85 (s, 3H); $^{13}\text{C}$ NMR (100 MHz, $\text{CDCl}_3$) $\delta$ 220.9, 212.8, 51.3, 47.8, 44.2, 42.2, 41.0, 37.1, 36.9, 35.9, 35.1, 35.0, 31.6, 26.3, 24.7, 22.6, 21.8, 20.5, 13.8.

**(3β,5β)-3-Hydroxyandrostane-17-one (4).** To a stirred solution of steroid **3** (1.65 g, 5.7 mmol) in dry THF (150 mL) was slowly added dropwise K-selectride (8 mL, 8 mmol, 1.0M in THF) at -78 °C over ~30 min. After 2 h, acetone (2 mL) was added. After 15 min, aqueous $\text{NH}_4\text{Cl}$ (20 mL) was added, the mixture was allowed to warm to 23 °C and stirring was continued for 1 h. The product was extracted into EtOAc (100 mL x 3). The combined extracts were dried over anhydrous $\text{Na}_2\text{SO}_4$, filtered and the solvent removed. The residue was purified by flash column chromatography (silica gel, eluted with 25% EtOAc and 25% $\text{CH}_2\text{Cl}_2$ in hexanes) to give steroid **4** (1.26 g, 76%): $^1\text{H}$ NMR (400 MHz, $\text{CDCl}_3$) $\delta$ 4.08 (s, 1H), 2.44-2.37 (m, 1H), 2.08-1.09 (m,

22H), 0.95 (s, 3H), 0.82 (s, 3H); $^{13}\text{C}$ NMR (100 MHz, $\text{CDCl}_3$) $\delta$ 221.6, 66.8, 51.5, 47.9, 40.0, 36.5, 35.9, 35.3, 35.2, 33.4, 31.7, 29.9, 27.8, 26.3, 25.2, 23.8, 21.8, 20.3, 13.8.

**(3β,5β)-((3-*tert*-Butyldimethylsilyl)oxy)-androstan-17-one (5).** To a stirred solution of steroid **4** (930 mg, 3.2 mmol) in DMF (10 mL) was added TBSCl (970 mg, 6.4 mmol) and imidazole (650 mg, 9.6 mmol) at 23 °C. After stirring overnight, aqueous $\text{NaHCO}_3$ was added and the product was extracted into EtOAc (100 mL). The extract was washed with brine (100 mL x 3), dried over anhydrous $\text{Na}_2\text{SO}_4$, filtered, the solvent removed and the residue purified by flash column chromatography (silica gel, eluted with 10% EtOAc in hexanes) to give steroid **5** (1.2 g, 93%): $^1\text{H}$ NMR (400 MHz, $\text{CDCl}_3$) $\delta$ 4.0 (s, 1H), 2.44-2.37 (m, 1H), 2.07-0.88 (m, 21H), 0.94 (s, 3H), 0.86 (s, 9H), 0.82 (s, 3H), -0.01 (s, 6H); $^{13}\text{C}$ NMR (100 MHz, $\text{CDCl}_3$) $\delta$ 221.5, 67.2, 51.6, 47.9, 40.2, 36.5, 35.9, 35.3, 35.2, 34.4, 31.8, 30.0, 28.6, 26.6, 25.8 (3 x C), 25.4, 23.9, 21.8, 20.4, 18.1, 13.8, -4.85, -4.87.

**(3β,5β,17β)-((3-*tert*-Butyldimethylsilyl)oxy)-androstan-17-ol (6).** To a stirred solution of steroid **5** (1.2 g, 2.96 mmol) in EtOH (50 mL) was added $\text{NaBH}_4$ (230 mg, 6 mmol) at 23 °C. After 2 h, aqueous $\text{NaHCO}_3$ (50 mL) was added and stirring continued for 15 min. Most of the EtOH was removed under reduced pressure and the remaining aqueous solution was extracted with EtOAc (100 mL x 2). The combined organic layers were dried over anhydrous $\text{Na}_2\text{SO}_4$, filtered, the solvent removed and the residue was purified by flash column chromatography (silica gel, eluted with 10-20% EtOAc in hexanes) to give steroid **6** 1.06 g, 88%): $^1\text{H}$ NMR (400 MHz, $\text{CDCl}_3$) $\delta$ 4.02 (s, 1H), 3.65-3.60 (m, 1H), 2.07-1.90 (m, 1H), 1.86-0.94 (m, 22H), 0.94 (s,

3H), 0.88 (s, 9H), 0.72 (s, 3H), 0.01 (s, 6H); $^{13}\text{C}$ NMR (100 MHz, $\text{CDCl}_3$) $\delta$ 82.1, 67.4, 51.2, 43.1, 40.2, 37.0, 36.6, 35.8, 35.1, 34.4, 30.6, 30.0, 28.6, 26.8, 26.1, 25.8 (3 x C), 24.0, 23.4, 20.7, 18.1, 11.1, -4.8, -4.9.

**(3β,5β,17β)-17-((4-Bromobenzyl)oxy)-3-((*tert*-butyldimethylsilyl)oxy)-androstane (7).** To a stirred solution of steroid **6** (1.06 g, 2.6 mmol) in THF (20 mL) was added KH (1.0 g, 30% in mineral oil, 7.5 mmol). The mixture was refluxed for 30 min. 4-Bromobenzyl bromide (1.3 g, 5.2 mmol) in THF (10 mL) was added and the reaction was refluxed for 16 h. After cooling, water was added and the product extracted into EtOAc (100 mL x 2). The combined extracts were dried over anhydrous $\text{Na}_2\text{SO}_4$, filtered, the solvent removed and the residue purified by flash column chromatography (silica gel, eluted with 2-5% EtOAc in hexanes) to give steroid **7** (1.4 g, 92%): $^1\text{H}$ NMR (400 MHz, $\text{CDCl}_3$) $\delta$ 7.46 (d, $J$ = 8.2 Hz, 2H), 7.21 (d, $J$ = 8.2 Hz, 2H), 4.48 (s, 2H), 4.02 (s, 1H), 3.39-3.35 (m, 1H), 1.93-0.94 (m, 22H), 0.94 (s, 3H), 0.88 (s, 9H), 0.80 (s, 3H), 0.01 (s, 6H); $^{13}\text{C}$ NMR (100 MHz, $\text{CDCl}_3$) $\delta$ 138.4, 131.3 (2 x C), 128.9 (2 x C), 121.0, 88.5, 70.8, 67.4, 51.4, 43.3, 40.2, 38.3, 36.6, 35.5, 35.1, 34.4, 30.0, 28.6, 28.0, 26.8, 26.0, 25.9 (3 x C), 24.0, 23.4, 20.8, 18.1, 11.9, -4.8, -4.9.

**1-(4-(((3 $\beta$ -((*tert*-Butyldimethylsilyl)oxy)-5 $\beta$ -androst-17 $\beta$ -yl)oxy)methyl)phenyl)-**

**2,2,2-trifluoroethan-1-one (8).** To a stirred solution of steroid **7** (1.4 g, 2.4 mmol) in THF (50 mL) was slowly added *n*-BuLi (2.5 M in THF, 2 mL, 5 mmol) at $-78^\circ\text{C}$. After 45 min, 1-trifluoroacetyl piperidine (1.3 g, 7 mmol) was added, and the mixture was stirred at $-78^\circ\text{C}$ for 1 h. Aqueous $\text{NaHCO}_3$ (10 mL) was added at $-78^\circ\text{C}$ and the reaction was slowly warmed to $23^\circ\text{C}$.

°C. The product was extracted into EtOAc ($2 \times 100$ mL). The combined extracts were dried, filtered, solvent removed and the residue purified by flash column chromatography (silica gel, eluted with 5-10% EtOAc in hexanes) to give steroid **8** (1.32 g, 93%): $^1\text{H}$ NMR (400 MHz, $\text{CDCl}_3$) $\delta$ 8.05 (d, $J = 8.2$ Hz, 2H), 7.52 (d, $J = 8.2$ Hz, 2H), 4.62 (s, 2H), 4.02 (s, 1H), 3.44-3.39 (m, 1H), 2.04-0.97 (m, 22H), 0.95 (s, 3H), 0.88 (s, 9H), 0.83 (s, 3H), 0.01 (s, 6H); $^{13}\text{C}$ NMR (100 MHz, $\text{CDCl}_3$) $\delta$ 180.3 (q, $J = 34.4$ Hz), 148.1, 130.21, 130.19, 128.8, 127.2 (2 x C), 121.0 (q, $J = 291.4$ Hz), 89.1, 70.6, 67.4, 51.4, 43.3, 40.2, 38.3, 36.5, 35.5, 35.1, 34.4, 30.0, 28.6, 27.9, 26.8, 26.0, 25.8 (3 x C), 24.0, 23.4, 20.8, 18.1, 11.9, -4.8, -4.9.

**1-(4-(((3β-Hydroxy-5β-androst-17β-yl)oxy)methyl)phenyl)-2,2,2-trifluoroethan-1-one (YX81).** To steroid **8** (820 mg, 1.2 mmol) was added THF (10 mL) and 6 N HCl (10 mL) at 23 °C. After stirring for 16 h, water was added, and the product was extracted into $\text{CH}_2\text{Cl}_2$ (2 x 150 mL). The extracts were washed with aqueous $\text{NaHCO}_3$, dried over anhydrous $\text{Na}_2\text{SO}_4$, filtered, solvent removed and the residue purified by flash column chromatography (silica gel, eluted with 20% EtOAc in hexanes) to give **YX81** (550 mg, 92%): $^1\text{H}$ NMR (400 MHz, $\text{CDCl}_3$) $\delta$ 8.05 (d, $J = 7.8$ Hz, 2H), 7.50 (d, $J = 7.8$ Hz, 2H), 4.62 (s, 2H), 4.10 (s, 1H), 3.43-3.39 (m, 1H), 2.04-1.02 (m, 23H), 0.97 (s, 3H), 0.83 (s, 3H); $^{13}\text{C}$ NMR (100 MHz, $\text{CDCl}_3$) $\delta$ 180.4 (q, $J = 35.1$ Hz), 148.1, 130.21, 130.20, 128.8, 127.2 (2 x C), 121.0 (q, $J = 291.5$ Hz), 89.1, 70.6, 67.1, 51.3, 43.3, 39.9, 38.2, 36.5, 35.4, 35.2, 33.5, 29.9, 27.9, 27.8, 26.5, 25.8, 23.9, 23.4, 20.7, 11.9.

**1-(4-(((3β-Sulfoxy-5β-androst-17β-yl)oxy)methyl)phenyl)-2,2,2-trifluoroethan-1-one, triethyl ammonium salt (YX82).** To a solution of **YX81** (240 mg, 0.54 mmol) in pyridine (5

mL) was added SO<sub>3</sub>-Me<sub>3</sub>N (200 mg, 1.4 mmol) at 23 °C and the reaction was stirred for 20 h. Pyridine was removed under reduced pressure and the residue was purified by flash column chromatography (silica gel, eluted with CH<sub>2</sub>Cl<sub>2</sub>/MeOH/Et<sub>3</sub>N = 100/10/1 (v/v/v)) to give **YX82** (318 mg, 89%): <sup>1</sup>H NMR (400 MHz, CDCl<sub>3</sub>) δ 7.98 (d, *J* = 7.8 Hz, 2H), 7.47 (d, *J* = 7.8 Hz, 2H), 4.66 (s, 1H), 4.56 (s, 2H), 3.38 (t, *J* = 7.8 Hz, 1H), 3.14 (q, *J* = 7.4 Hz, 6H), 1.97-0.93 (m, 23H), 1.34 (t, *J* = 7.4 Hz, 9H), 0.88 (s, 3H), 0.76 (s, 3H); <sup>13</sup>C NMR (100 MHz, CDCl<sub>3</sub>) δ 180.6 (q, *J* = 35.1 Hz), 148.2, 130.14, 130.11, 128.6, 127.2 (2 x C), 121.0 (q, *J* = 291.4 Hz), 89.0, 75.3, 70.6, 51.2, 46.5 (3 x C), 43.3, 40.1, 38.1, 36.8, 35.4, 34.7, 31.3, 30.4, 27.9, 26.3, 25.7, 23.7, 23.3, 20.7, 11.8, 8.7 (3 x C).

### Synthesis of YX83 and YX84

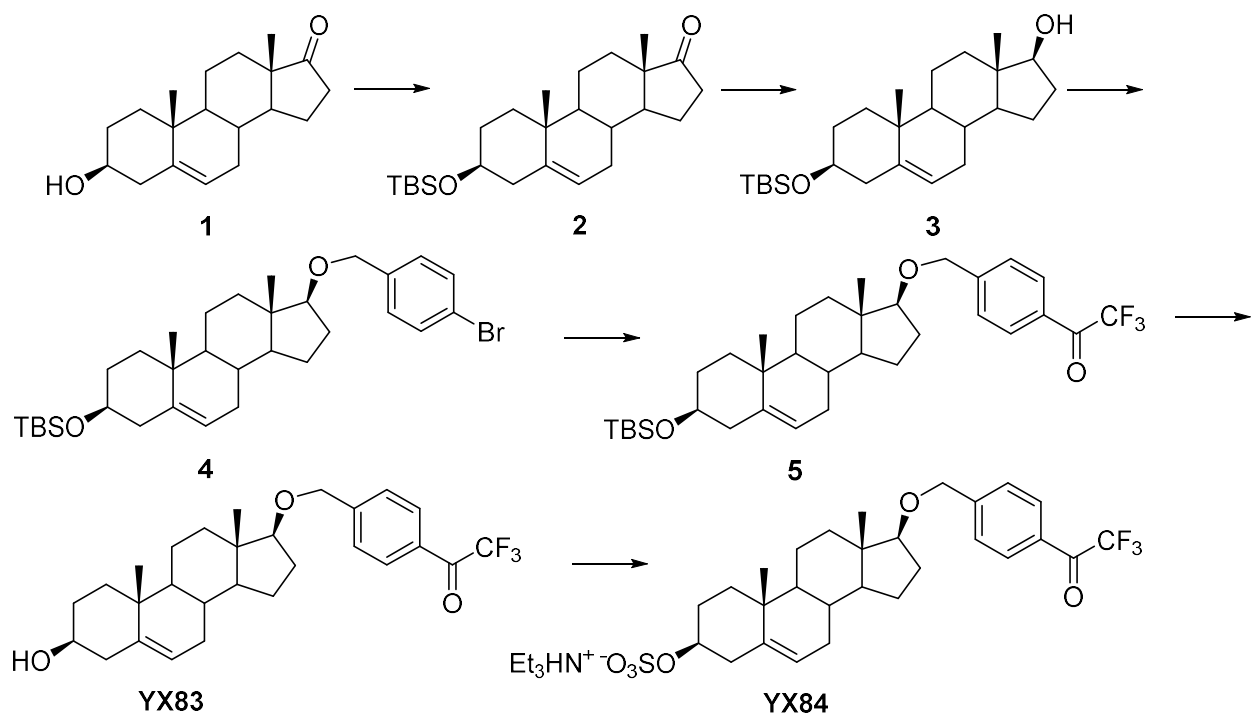

**3β-(*tert*-Butyldimethylsilyloxy)-androst-5-en-17-one (2).** To a stirred solution of dehydroepiandrosterone (**1**, 1.0 g, 3.5 mmol) in DMF (10 mL) was added TBSCl (1.1 g, 7 mmol) and imidazole (710 mg, 10.5 mmol) at 23 °C. After stirring overnight, aqueous NaHCO<sub>3</sub> was added and the product was extracted into EtOAc (100 ml). The EtOAc was washed with brine (100ml x 3), dried over anhydrous Na<sub>2</sub>SO<sub>4</sub>, filtered, solvent removed and the residue purified by flash column chromatography (silica gel, eluted with 5% EtOAc in hexanes) to give steroid **2** (1.4 g, ~100 %): <sup>1</sup>H NMR (400 MHz, CDCl<sub>3</sub>) $\delta$ 5.34-5.32 (m, 1H), 3.51-3.43 (m, 1H), 2.47-2.41 (m, 1H), 2.29-0.94 (m, 18H), 1.01 (s, 3H), 0.87 (s, 12H), 0.04 (s, 6H); <sup>13</sup>C NMR (100 MHz,

CDCl<sub>3</sub>) $\delta$ 221.1, 141.7, 120.4, 72.4, 51.8, 50.3, 47.5, 42.8, 37.3, 36.7, 35.8, 32.0, 31.5, 31.4, 30.8, 25.9 (3 x C), 21.9, 20.3, 19.4, 18.2, 13.5, -4.6 (2 x C).

**(3β,17β)-((*tert*-Butyldimethylsilyl)oxy)-androst-5-en-17-ol (3).** To a stirred solution of steroid **2** (1.4 g, 3.5 mmol) in EtOH (50 mL) was added NaBH<sub>4</sub> (270 mg, 7 mmol) at 23 °C. After 2 h, aqueous NaHCO<sub>3</sub> (50 mL) was added and stirring continued for 15 min. Most of EtOH was removed under reduced pressure and the remaining aqueous solution was extracted into EtOAc (100 mL x 2). The combined extracts were dried over anhydrous Na<sub>2</sub>SO<sub>4</sub>, filtered, solvent removed and the residue purified by flash column chromatography (silica gel, eluted with 10-20% EtOAc in hexanes) to give steroid **3** (1.2 g, 86%): <sup>1</sup>H NMR (400 MHz, CDCl<sub>3</sub>) $\delta$ 5.31-5.30 (m, 1H), 3.65-3.61 (m, 1H), 3.48-3.46 (m, 1H), 2.26-0.92 (m, 20H), 1.00 (s, 3H), 0.88 (s, 9H), 0.75 (s, 3H), 0.05 (s, 6H); <sup>13</sup>C NMR (100 MHz, CDCl<sub>3</sub>) $\delta$ 82.1, 67.4, 51.2, 43.1, 40.2, 37.0, 36.6, 35.8, 35.1, 34.4, 30.6, 30.0, 28.6, 26.8, 26.1, 25.8 (3 x C), 24.0, 23.4, 20.7, 18.1, 11.1, -4.8, -4.9.

**(3β,17β)-17-((4-Bromobenzyl)oxy)-3-((*tert*-butyldimethylsilyl)oxy)-androst-5-ene (4).** To a stirred solution of steroid **3** (1.2 g, 3 mmol) in THF (20 mL) was added KH (1.0 g, 30% in mineral oil, 7.5 mmol). The mixture was refluxed for 30 min. 4-Bromobenzyl bromide (1.5 g, 6 mmol) in THF (10 mL) was added and the reaction was refluxed for 16 h. After cooling, water was added and the product was extracted into EtOAc (100 mL x 2). The combined extracts were dried over anhydrous Na<sub>2</sub>SO<sub>4</sub>, filtered, solvent removed and the residue purified by flash column chromatography (silica gel, eluted with 2-5% EtOAc in hexanes) to give steroid **4** (1.6 g, 93%): <sup>1</sup>H NMR (400 MHz, CDCl<sub>3</sub>) $\delta$ 7.46 (d, *J* = 7.4 Hz, 2H), 7.22 (d, *J* = 7.4 Hz, 2H), 5.32-5.31 (m,

1H), 4.49 (s, 2H), 3.54-3.46 (m, 1H), 3.41-3.37 (m, 1H), 2.31-0.92 (m, 19H), 1.02 (s, 3H), 0.90 (s, 9H), 0.84 (s, 3H), 0.07 (s, 6H); <sup>13</sup>C NMR (100 MHz, CDCl<sub>3</sub>) δ 141.6, 138.4, 131.6, 131.3, 129.4, 128.9, 121.0, 120.9, 88.4, 72.6, 70.9, 51.5, 50.3, 42.9, 42.8, 37.8, 37.4, 36.7, 32.1, 31.7, 31.5, 27.9, 26.0 (3 x C), 23.5, 20.7, 19.5, 18.3, 11.7, -4.6 (2 x C).

**1-(4-(((3β-((*tert*-Butyldimethylsilyl)oxy)-androst-5-ene-17β-yl)oxy)methyl)phenyl)-2,2,2-trifluoroethan-1-one (5).** To a stirred solution of steroid **4** (1.6 g, 2.8 mmol) in THF (50 mL) was slowly added *n*-BuLi (2.5 M in THF, 2 mL, 5 mmol) at -78 °C. After 45 min, 1-trifluoroacetyl piperidine (1.3 g, 7 mmol) was added and the mixture was stirred at -78 °C for 1 h. Aqueous NaHCO<sub>3</sub> (10 mL) was added at -78 °C and the reaction was slowly warmed to 23 °C. The product was extracted into EtOAc (2 x 100 mL). The combined extracts were dried over anhydrous Na<sub>2</sub>SO<sub>4</sub>, filtered, solvent removed and the residue purified by flash column chromatography (silica gel, eluted with 5-10% EtOAc in hexanes) to give steroid **5** (1.62 g, 97%): <sup>1</sup>H NMR (400 MHz, CDCl<sub>3</sub>) δ 8.06 (d, *J* = 8.2 Hz, 2H), 7.52 (d, *J* = 8.2 Hz, 2H), 5.32-5.31 (m, 1H), 4.63 (s, 2H), 3.51-3.41 (m, 2H), 2.27-0.93 (m, 20H), 1.02 (s, 3H), 0.89 (s, 9H), 0.86 (s, 3H), 0.06 (s, 6H); <sup>13</sup>C NMR (100 MHz, CDCl<sub>3</sub>) δ 180.3 (q, *J* = 35.1 Hz), 148.1, 141.6, 130.22, 130.20, 128.8, 127.2 (2 x C), 121.0 (q, *J* = 291.4 Hz), 120.8, 89.0, 72.5, 70.7, 51.5, 50.3, 43.0, 42.8, 37.8, 37.4, 36.6, 32.0, 31.7, 31.5, 27.8, 25.9 (3 x C), 23.5, 20.7, 19.5, 18.3, 11.7, -4.6 (2 x C).

**1-(4-(((3β-Hydroxy-androst-5-ene-17β-yl)oxy)methyl)phenyl)-2,2,2-trifluoroethan-1-one (YX83).** To steroid **5** (620 mg, 1 mmol) was added THF (10 mL) and 6 N HCl (10 mL) at 23 °C.

After 16 h, water was added and the product was extracted into CH<sub>2</sub>Cl<sub>2</sub> (2 x 150 mL). The combined organic extracts were washed with aqueous NaHCO<sub>3</sub>, dried over anhydrous Na<sub>2</sub>SO<sub>4</sub>, filtered, solvent removed and the residue purified by flash column chromatography (silica gel, eluted with 20% EtOAc in hexanes) to give **YX83** (400 mg, 84%): <sup>1</sup>H NMR (400 MHz, CDCl<sub>3</sub>) δ 8.04 (d, *J* = 8.1 Hz, 2H), 7.51 (d, *J* = 8.1 Hz, 2H), 5.33-5.32 (m, 1H), 4.62 (s, 2H), 3.53-3.40 (m, 2H), 2.29-0.92 (m, 20H), 1.00 (s, 3H), 0.85 (s, 3H); <sup>13</sup>C NMR (100 MHz, CDCl<sub>3</sub>) δ 180.3 (q, *J* = 34.3 Hz), 148.1, 140.9, 130.21, 130.19, 128.8, 127.2 (2 x C), 121.3, 121.0 (q, *J* = 291.5 Hz), 89.0, 71.6, 70.7, 51.5, 50.2, 42.9, 42.2, 37.8, 37.3, 36.5, 31.7, 31.6, 31.4, 27.8, 23.4, 20.7, 19.4, 11.7.

**1-(4-(((3β-Sulfoxy-androst-5-ene-17β-yl)oxy)methyl)phenyl)-2,2,2-trifluoroethan-1-one, triethyl ammonium salt (YX84).** To a stirred solution of **YX83** (240 mg, 0.54 mmol) in pyridine (5 mL) was added SO<sub>3</sub>-Me<sub>3</sub>N (200 mg, 1.4 mmol) at 23 °C. The reaction was stirred at room temperature for 20 h. Pyridine was removed under reduced pressure and the residue was purified by flash column chromatography (silica gel, eluted with CH<sub>2</sub>Cl<sub>2</sub>/MeOH/Et<sub>3</sub>N = 100/10/1 (v/v/v)) to give **YX84** (260 mg, 79%): <sup>1</sup>H NMR (400 MHz, CDCl<sub>3</sub>) δ 7.98 (d, *J* = 8.2 Hz, 2H), 7.46 (d, *J* = 8.2 Hz, 2H), 5.29-5.28 (m, 1H), 4.56 (s, 2H), 3.38 (t, *J* = 7.8 Hz, 1H), 3.14 (q, *J* = 7.4 Hz, 6H), 2.52-2.49 (m, 1H), 2.33-2.27 (m, 1H), 2.05-0.83 (m, 19H), 1.31 (t, *J* = 7.4 Hz, 9H), 0.94 (s, 3H), 0.78 (s, 3H); <sup>13</sup>C NMR (100 MHz, CDCl<sub>3</sub>) δ 180.3 (q, *J* = 35.1 Hz), 148.1, 140.3, 130.14, 130.12, 128.6, 127.2 (2 x C), 121.8, 121.0 (q, *J* = 291.4 Hz), 88.9, 78.0, 70.6, 51.3, 50.1, 46.5 (3 x C), 42.9, 39.3, 37.7, 37.1, 36.5, 31.6, 31.4, 28.9, 27.8, 23.4, 20.6, 19.3, 11.7, 8.8 (3 x C).

### Synthesis of MQ364 and MQ384

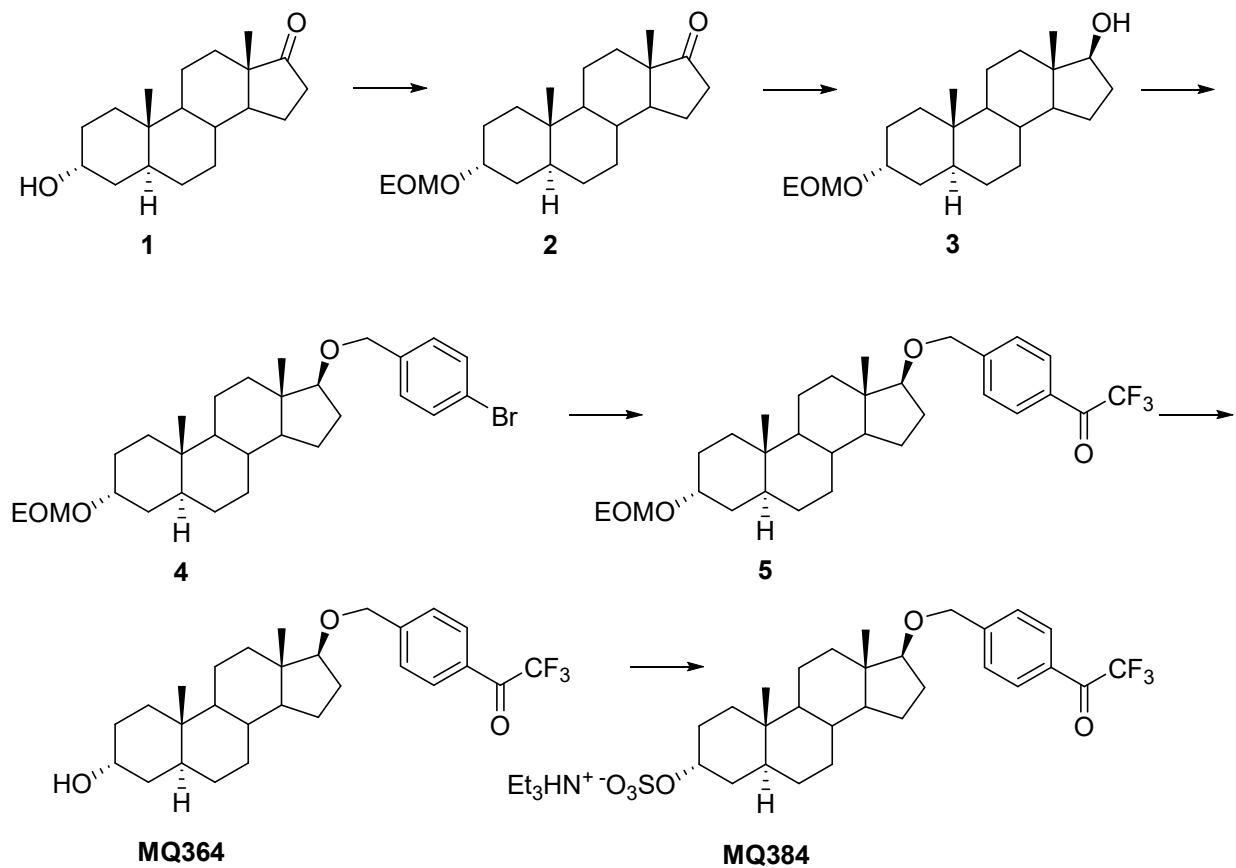

**(3α,5α)-3-(Ethoxymethoxy)-androstan-17-one (2).** To a stirred solution of androsterone (**1**, 3 g, 10.34 mmol) in $\text{CH}_2\text{Cl}_2$ (40 mL) was added chloromethyl ethyl ether (1.86 mL, 20 mmol) and $(i\text{-Pr})_2\text{NEt}$ (4.2 mL, 30 mmol) at 23 °C for 16 h. Solvent was removed and the residue was purified by flash column chromatography (silica gel, eluted with 15% EtOAc in hexanes) to give steroid **2** (3.54 g, 98%): $^1\text{H}$ NMR (400 MHz, $\text{CDCl}_3$) $\delta$ 4.56 (q, $J = 7.0$ Hz, 2H), 3.69 (s, 1H), 3.47 (q, $J = 7.0$ Hz, 2H), 2.30 (dd, $J = 9.5$ Hz, 10.1 Hz, 1H), 1.95-0.63 (m, 21H), 1.06 (t, $J = 7.0$ Hz, 3H), 0.70 (s, 3H), 0.67 (s, 3H); $^{13}\text{C}$ NMR (100 MHz, $\text{CDCl}_3$) $\delta$ 220.5, 92.7, 71.0, 62.5, 54.1, 51.1, 47.3, 39.3, 35.6, 35.4, 34.6, 33.2, 32.4, 31.2, 30.5, 27.9, 25.9, 21.4, 19.7, 14.9, 13.4, 11.0.

**(3α,5α,17β)-3-(Ethoxymethoxy)-androstane-17-ol (3).** To a stirred solution of steroid **2** in EtOH (50 mL) was added NaBH<sub>4</sub> (780 mg, 20 mmol) at 23 °C. After 1 h, aqueous NH<sub>4</sub>Cl was added and stirring continued for 15 min. EtOH was removed and water (50 mL) was added to the remaining aqueous solution. The product was extracted into EtOAc (150 mL x 2). The combined extracts were dried over anhydrous Na<sub>2</sub>SO<sub>4</sub>, filtered, solvent removed and the residue purified by flash column chromatography (silica gel, eluted with 35% EtOAc in hexanes) to give steroid **3** (2.9 g, 82%): <sup>1</sup>H NMR (400 MHz, CDCl<sub>3</sub>) $\delta$ 4.69 (q, $J$ = 7.0 Hz, 2H), 3.81 (s, 1H), 3.60-3.54 (m, 3H), 2.00-0.69 (m, 23H), 1.19 (t, $J$ = 7.0 Hz, 3H), 0.76 (s, 3H), 0.69 (s, 3H); <sup>13</sup>C NMR (100 MHz, CDCl<sub>3</sub>) $\delta$ 92.9, 81.7, 71.4, 62.9, 54.3, 51.0, 42.9, 39.7, 36.7, 35.8, 35.4, 33.5, 32.7, 31.4, 30.3, 28.4, 26.2, 23.2, 20.2, 15.1, 11.3, 11.1.

**(3α,5α,17β)-17-((4-Bromobenzyl)oxy)-3-(ethoxymethoxy)-androstane (4).** To a stirred solution of steroid **3** (500 mg, 1.43 mmol) in THF (30 mL) was added KH (600 mg, 30% in mineral oil, 4.5 mmol). The mixture was refluxed for 30 min. 4-Bromobenzyl bromide (1.08 g, 4.3 mmol) in THF (10 mL) was added and reflux continued for 16 h. After cooling, water was added and the product was extracted into EtOAc (100 mL x 2). The combined extracts were dried over anhydrous Na<sub>2</sub>SO<sub>4</sub>, filtered, solvent removed and the residue purified by flash column chromatography (silica gel, eluted with 2-5% EtOAc in hexanes) to give steroid **4** (740 mg, 100%): <sup>1</sup>H NMR (400 MHz, CDCl<sub>3</sub>) $\delta$ 7.44 (d, $J$ = 8.1 Hz, 2H), 7.20 (d, $J$ = 8.1 Hz, 2H), 4.72 (q, $J$ = 7.0 Hz, 2H), 4.46 (s, 2H), 3.84 (s, 1H), 3.63 (q, $J$ = 7.0 Hz, 2H), 3.38 (t, $J$ = 8.2 Hz, 1H), 1.97-0.73 (m, 22H), 1.22 (t, $J$ = 7.0 Hz, 3H), 0.80 (s, 3H), 0.79 (s, 3H); <sup>13</sup>C NMR (100 MHz, CDCl<sub>3</sub>) $\delta$ 138.3, 131.2 (2 x C), 128.7 (2 x C), 120.9, 93.0, 88.4, 71.4, 70.7, 62.8, 54.3, 51.1, 43.0, 39.6, 37.9, 35.8, 35.1, 33.4, 32.7, 31.4, 28.4, 27.8, 26.2, 23.2, 20.3, 15.1, 11.8, 11.3.

**1-(4-(((3α-(Ethoxymethoxy)-5α-androstan-17β-yl)oxy)methyl)phenyl)-2,2,2-trifluoroethan-1-one (5).** To a stirred solution of steroid **4** (450 mg, 0.87 mmol) in THF (20 mL) was slowly added *n*-BuLi (2.5 M in THF, 1.04 mL, 2.6 mmol) at -78 °C. After 45 min, 1-trifluroacetyl piperidine (725 mg, 4 mmol) was added and stirring was continued at -78 °C for 1 h. Aqueous NaHCO<sub>3</sub> (5 mL) was added at -78 °C and the reaction was slowly warmed to 23 °C. The product was extracted into EtOAc (150 mL x 2). The combined extracts were dried over anhydrous Na<sub>2</sub>SO<sub>4</sub>, filtered, solvent removed and the residue purified by flash column chromatography (silica gel, eluted with 15% EtOAc in hexanes) to give steroid **5** (209 mg, 45%): <sup>1</sup>H NMR (400 MHz, CDCl<sub>3</sub>) $\delta$ 8.02 (d, *J* = 8.2 Hz, 2H), 7.50 (d, *J* = 8.2 Hz, 2H), 4.70 (q, *J* = 7.0 Hz, 2H), 4.60 (s, 2H), 3.82 (s, 1H), 3.61 (q, *J* = 7.0 Hz, 2H), 3.42 (t, *J* = 8.2 Hz, 1H), 1.93-0.73 (m, 22H), 1.20 (t, *J* = 7.0 Hz, 3H), 0.81 (s, 3H), 0.78 (s, 3H); <sup>13</sup>C NMR (100 MHz, CDCl<sub>3</sub>) $\delta$ 180.2 (q, *J* = 35.1 Hz), 148.1, 130.2 (2 x C), 128.7, 127.2 (2 x C), 118.4 (q, *J* = 282.4 Hz), 93.1, 89.1, 71.5, 70.6, 62.9, 54.4, 51.2, 43.2, 39.7, 38.0, 35.9, 35.3, 33.6, 32.8, 31.5, 28.5, 27.8, 26.3, 23.4, 20.4, 15.1, 11.9, 11.4.

**1-(4-(((3α-Hydroxy-5α-androst-17β-yl)oxy)methyl)phenyl)-2,2,2-trifluoroethan-1-one (MQ364).** Steroid **5** (350 mg, 0.57 mmol) was stirred in THF/ 6 N HCl (10 mL/10 mL) at 23 °C. After 2 h, water was added and the product was extracted into CH<sub>2</sub>Cl<sub>2</sub> (150 mL x 2). The combined extracts were washed with aqueous NaHCO<sub>3</sub>, dried over anhydrous Na<sub>2</sub>SO<sub>4</sub>, filtered, solvent removed and the residue purified by flash column chromatography (silica gel, eluted with 25–35% EtOAc in hexanes) to give **MQ364** (157 mg, 84%): <sup>1</sup>H NMR (400 MHz, CDCl<sub>3</sub>) $\delta$ 8.03 (d, *J* = 8.1 Hz, 2H), 7.50 (d, *J* = 8.1 Hz, 2H), 4.61 (s, 2H), 4.01 (s, 1H), 3.43 (t, *J* = 8.1 Hz, 1H),

2.03-0.73 (m, 23H), 0.82 (s, 3H), 0.77 (s, 3H); $^{13}\text{C}$ NMR (100 MHz, $\text{CDCl}_3$) $\delta$ 180.3 (q, $J = 36.1$ Hz), 148.1, 130.14 (2 x C), 130.12, 127.1 (2 x C), 118.1 (q, $J = 281.5$ Hz), 89.1, 70.6, 66.4, 54.4, 51.2, 43.1, 39.0, 37.9, 36.1, 35.8, 35.2, 32.1, 31.5, 28.9, 28.3, 27.8, 23.3, 20.4, 11.8, 11.1.

**1-(4-(((3 $\alpha$ -Sulfoxy-5 $\alpha$ -androst-17 $\beta$ -yl)oxy)methyl)phenyl)-2,2,2-trifluoroethan-1-one,**

**triethyl ammonium salt (MQ384).** To a stirred solution of **MQ364** (36 mg, 0.0756 mmol) in pyridine (3 mL) was added $\text{SO}_3\text{-Me}_3\text{N}$ (53 mg, 0.378 mmol) at 23 °C. The mixture was stirred at 23 °C for 20 h. Pyridine was removed under reduced pressure and the residue was purified by flash column chromatography (silica gel, eluted with $\text{CH}_2\text{Cl}_2/\text{MeOH}/\text{Et}_3\text{N} = 100/10/1$ (v/v/v)) to give **MQ384** (34 mg, 68%): $^1\text{H}$ NMR (400 MHz, $\text{CDCl}_3$) $\delta$ 8.03 (d, $J = 8.2$ Hz, 2H), 7.50 (d, $J = 8.2$ Hz, 2H), 4.64 (s, 1H), 4.61 (s, 2H), 3.45 (t, $J = 8.1$ Hz, 1H), 3.14 (q, $J = 7.4$ Hz, 6H), 1.99-0.77 (m, 23H), 1.39 (t, $J = 7.4$, 9H), 0.81 (s, 3H), 0.77 (s, 3H); $^{13}\text{C}$ NMR (100 MHz, $\text{CDCl}_3$) $\delta$ 180.3 (q, $J = 35.3$ Hz), 130.1 (2 x C), 128.6, 127.1 (2 x C), 118.1 (q, $J = 291.5$ Hz), 89.0, 74.8, 70.5, 54.2, 51.2, 46.1 (3 x C), 43.1, 39.2, 38.0, 35.6, 35.2, 33.6, 32.5, 31.4, 28.1, 27.8, 26.7, 23.3, 20.3, 11.8, 11.4, 8.6 (3 x C).
